# Conserved reduction of m^6^A marks during aging and neurodegeneration is linked to altered translation of synaptic transcripts

**DOI:** 10.1101/2022.06.08.495100

**Authors:** Ricardo Castro-Hernández, Tea Berulava, Maria Metelova, Robert Epple, Tonatiuh Peña Centeno, M Sadman Sakib, Susanne Burkhart, Momchil Ninov, Katherine E. Bohnsack, Markus T. Bohnsack, Ivana Delalle, Andre Fischer

**Author notes:** equal contribution (Ricardo Castro-Hernández, Tea Berulava).

## Abstract

N^6^-methyladenosine (m^6^A) plays diverse roles in the regulation of mRNA metabolism. In the mammalian brain it has been linked to developmental processes and memory function. However, the precise role of m^6^A in the context synaptic plasticity and especially during impaired cognition are not fully understood. Here, we describe the mouse and human brain m^6^A epi-transcriptome in a tissue-specific manner. We furthermore show that m6A levels undergo a massive decrease across mouse brain regions as a consequence of aging. In addition, Alzheimer's disease in humans correlates with decreased N^6^-methylation in a similar population of transcripts that are linked to synaptic function and localized to synapses, such as the calcium/calmodulin-dependent kinase II (CaMKII). We furthermore show that reduced m6A levels impair synaptic protein-synthesis of CAMKII. Our results suggest that m^6^A-RNA-methylation is an important mechanism to control synaptic protein synthesis which is affected early in cognitive diseases.

**Significance statement:** The addition of N^6^-methyladenosine (m^6^A) to RNA plays a role in various cellular processes and its de-regulation has been linked to several devastating diseases. The precise role of m^6^A RNA-methylation in the adult brain is, however, not well understood. In our study, we describe the genome-wide m^6^A epi-transcriptome in the healthy and diseased brains of mice and humans. Our data demonstrate that a substantial amount of m^6^A transcripts are conserved. These transcripts are linked to the regulation of synaptic processes and are localized to synapses. In the diseases brain we detect RNA hypomethylation across multiple transcripts in all investigated brain regions and across species. At the mechanistic level we find that reduced m^6^A levels specifically impairs synaptic protein-synthesis.

## Introduction

Decades after it was first described, the posttranscriptional modification and labeling of specific nucleotides in mRNA has become the target of intense research interest in recent years ^1^. The most abundant of these marks, N^6^-methyladenosine (m^6^A), has been at the forefront of research in multiple fields of biology due to its dynamic nature as well as the broad range of molecular consequences for m^6^A-labeled transcripts, giving rise to the field of epitranscriptomics ^2 3 4^.

The deposition of m^6^A methylation marks on targeted mRNAs is made possible by the activity of a m^6^A methylation complex formed by the methyltransferases (METTL) METTL3 and METTL14 with the adaptor protein WTAP ^5 6 7^. M^6^A sites can be found across the entire transcript but often concentrate close to the stop codon and in the 3’UTR, falling in the motif consensus sequence DRACH (where D= A,T or G, R= A or G, and H= A, T or C ^2 3 8^. These methylation marks can in turn be removed by demethylases like Fat Mass And Obesity-Associated Protein (FTO) and Alpha-Ketoglutarate-Dependent Dioxygenase AlkB Homolog 5 (ALKBH5), making the regulation of m^6^A levels a complex and highly dynamic process ^9 10 11^. m^6^A labeled transcripts are recognized by a wide array of reader proteins and thus m^6^A RNA-methylation can affect a broad array of processes associated with mRNA metabolism, including nuclear export, transport, degradation and translation ^12 13 14 15 7 16^.

These properties have brought the epitranscriptome forward as a key component of the intricate regulatory networks that rule complex physiological and pathological processes such as development or the pathogenesis of cancer ^17 18 19 20 21^. While m^6^A marks are widespread and highly dynamic they are highest in the adult mammalian brain, and in recent years research has focused on deciphering its role in the regulation of brain function ^3 22^. Thus, m6A-dependent regulation in the adult brain has been linked to memory consolidation, learning and injury recovery ^23 24 25 26 27 28 29^.

More recently researchers started to investigate m^6^A levels in neurodegenerative diseases ^30 21 31 32 33^. While changes in m^6^A levels have been observed, the magnitude, depth, directionality and functional consequences of these changes are still a matter of contention in the field ^34 30 35 36 33^.

In this study we analyze the m^6^A epitranscriptome across multiple regions of the adult mouse and human brain. We find a remarkable conservation of methylation marks between mouse and human related to transcripts that are linked to synapse function, while other processes such as gene-expression control appear to be species-specific. Differential methylation analyses of cognitively impaired aged mice and human AD patients revealed a stark m^6^A decrease within transcripts involved in synapse function in multiple brain subregions. The genes affected by m^6^A changes converge on multiple synaptic plasticity-associated pathways in both aging and AD, among them CAMKII, a key regulator of synaptic signaling. Finally, we show that reducing m^6^A levels within *CamkII* transcripts results in impaired synaptic synthesis of the corresponding protein, suggesting that loss of m^6^A marks on transcripts associated with synaptic function and plasticity, is an early event in cognitive diseases.

## Results

### The m^6^A landscape in the adult mouse brain

Recent studies have implicated N-6-methyladenosine modifications in mRNA with learning and synaptic plasticity ^20 22^. Our knowledge about the transcriptome-wide distribution of m^6^A marks as well as the downstream molecular mechanisms involved in the regulation of neuronal plasticity is however still limited. Thus, we started our analysis by characterizing the landscape of m^6^A modifications in the healthy adult brain. The brains of ten (C57BL/6J) 3-month-old (young) wild type (WT) mice were extracted and dissected to obtain hippocampal subregions: CA1, CA3 and dentate gyrus (DG); as well as the anterior cingulate cortex (ACC, Figure 1A). meRIP-seq was performed on the mRNAs extracted from these samples to determine the subregion-specific epitranscriptome landscape in young adult mice.

**Figure 1.**
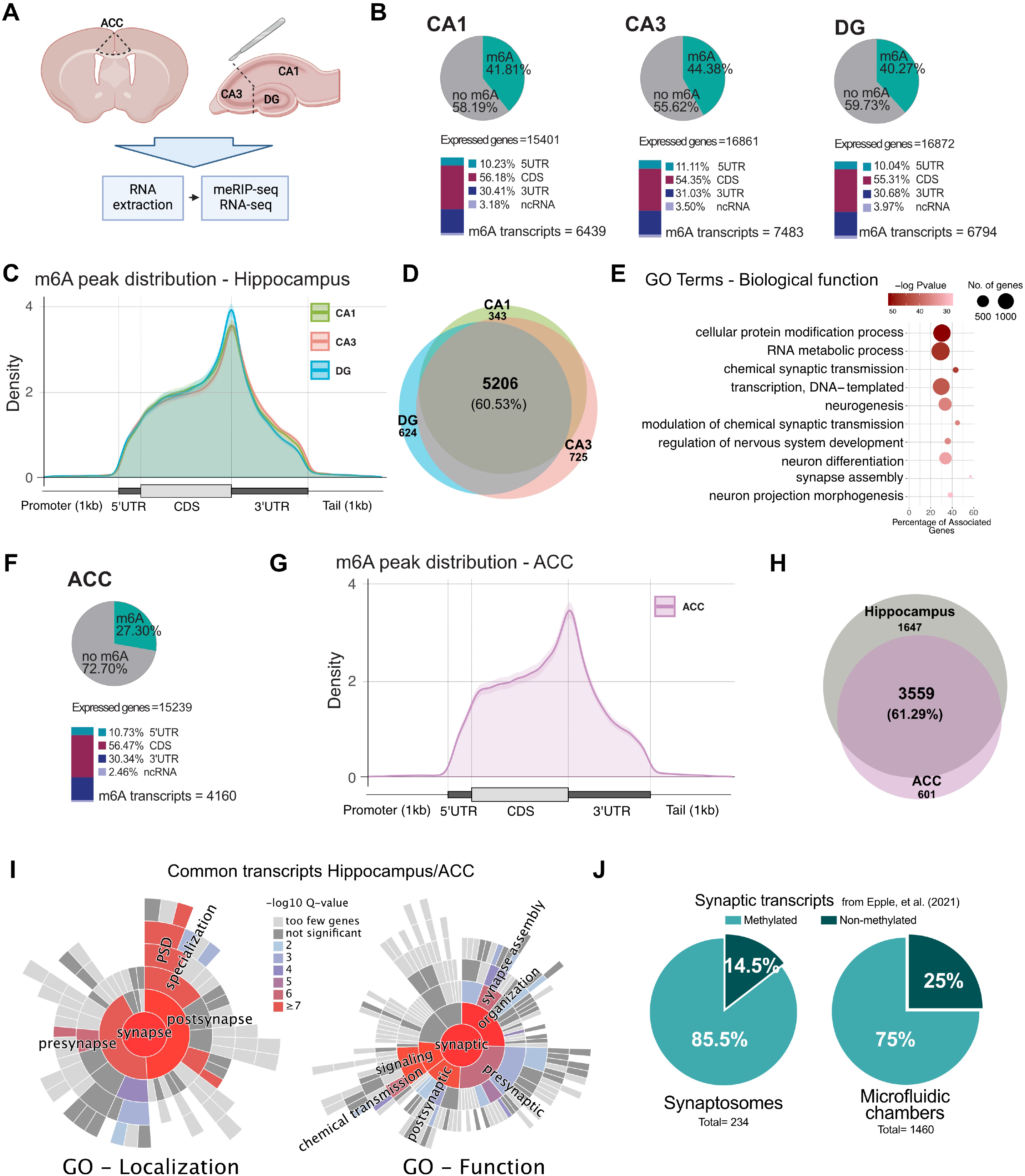
The m^6^A epitranscriptome in the adult mouse brain. **A**. Experimental scheme for the dissection of brain subregions for meRIP and RNA-seq experiments. **B**. Distribution of methylation across the transcriptome in the hippocampal subregions. Percentage of m^6^A calculated against the corresponding expressed genes in inputs; annotated peak regions calculated from total m^6^A peaks. **C**. Guitar plot showing the distribution of m^6^A peaks along mRNAs in hippocampal subregions. **D**. Overlap in methylated transcripts across hippocampal subregions. **E**. Enrichment of biological function Gene Ontology terms in commonly methylated hippocampal transcripts. **F**. Distribution of methylation across the transcriptome in the ACC. **G**. Guitar plot showing the distribution of m^6^A peaks along mRNA features in the ACC. **H**. Overlap between methylated transcripts in the Hippocampus (common transcripts from D) and the ACC. **I**. Enrichment of synapse-specific GO terms in commonly methylated transcripts between the hippocampus and ACC. **J**. Percentage of methylated transcripts from an independent synaptic mRNA dataset obtained from the hippocampus ^38^. ACC - anterior cingulate cortex, DG - dentate gyrus, 5UTR - 5’ untranslated region, 3UTR - 3’ untranslated region, CDS - coding sequence, ncRNA - non-coding RNA.

The analysis of methylated regions showed a large number of detected m^6^A peaks in hippocampal brain subregions, with 18270 peaks detected in the CA1, 20415 in the CA3 and 16686 in the DG (Figure S1A). A remarkable number of transcripts were detected carrying this methylation mark, ranging from 40.27% of the expressed genes in the DG to 42.81% and 44.38% in the CA1 and CA3 regions, respectively (Figure 1B, Suppl. Table 1). On average, every methylated transcript had 2.4-2.7 methylated regions per transcript containing m^6^A, depending on the hippocampal subregion (Figure S1B). Motif enrichment analyses of the detected m^6^A peaks showed a strong overrepresentation of the consensus m^6^A motif DRACH, showing that the meRIP-Seq had successfully enriched for m^6^A sites (Figure S1C). The detected m^6^A peaks follow a distribution along transcripts that corresponds with the well-described location of m^6^A sites, with enrichment in the vicinity of the stop codon and 3’UTR, as well as internal exons (Figure 1B,C). The population of methylated transcripts exhibited a large similarity, with about 60% of all transcripts with m^6^A in all subregions being common across them. However, despite the large overlap, a subset of transcripts appeared to be methylated in a subregion-specific manner, ranging from 6.18% in CA1 to 12.21% of the total in CA3 (Figure 1D, Suppl. Table 2).

The 5206 transcripts that are detected as being methylated in all hippocampal subregions showed a very significant enrichment for genes associated with neurogenesis and neural development, RNA metabolism, as well as synapse assembly and function (Figure 1E), supporting the notion that m^6^A acts as a crucial regulator of these processes in the adult brain. Subregion-specific transcripts showed an enrichment for more broad biological processes with no clear mechanism standing out (Figure S1D-F).

To further understand how the epitranscriptomic landscape varies across brain subregions, we performed the meRIP-seq analysis on ACC samples obtained from the same young mice. The mouse ACC showed a considerable but reduced methylation level, compared to the hippocampus. In the ACC 11816 m^6^A peaks were detected, corresponding to 4160 consistently methylated transcripts (2.83 peaks per transcript), which represented 27.3% of the expressed genes, highlighting the remarkable tissue specificity of RNA methylation (Figure 1F, Figure S1A,B).

Methylated transcripts in the ACC followed a similar distribution pattern along their sequence with enrichment in the CDS and 3’UTR (Figure 1F,G). Interestingly, the hippocampus and ACC shared a 61.29% of their methylated transcripts (Figure 1H, Suppl. Table 3). Commonly methylated transcripts across brain subregions showed a very strong enrichment in pathways associated with synaptic assembly, organization and signaling, as well as learning and memory, similar to what could be observed in ACC-specific mRNAs (Figure S2A,B). In contrast, hippocampus-specific transcripts have functions in the regulation of gene expression and RNA metabolism (Figure S2C).

Using SYNGO ^37^, an experimentally annotated database for synaptic location and function gene ontology (GO), we could confirm that commonly methylated transcripts between hippocampus and ACC are highly enriched for synaptically located proteins (Figure 1I). ACC-specific transcripts also display a very significant enrichment for synaptic proteins. In contrast, hippocampus-specific transcripts showed no significant synaptic enrichment (Figure S2D).

These data suggest that m^6^A transcripts might be specifically enriched at synapses, which is in agreement with previous data that detected m^6^A mRNAs in synaptosomes ^27^. To further explore this hypothesis, we made use of a recently published dataset containing a high-confidence hippocampal synaptic RNAome and compared it to our hippocampal epitranscriptome data. This synaptic RNA dataset was generated from purified synaptosomes of WT mice as well as primary neurons grown in microfluidic chambers to isolate their synaptic compartments, making it a robust resource of synaptically located RNAs ^38^ (Figure S2E,F). In both datasets, we observed a strong enrichment of methylated transcripts in synaptically located mRNAs with more than 70% of the synaptosome and 64% of the microfluid chamber transcriptome having at least one m^6^A peak (Figure 1J, Suppl. Table 4). These results go in accordance with previous reports describing the epitranscriptome as largely constant across brain regions, with comparatively small tissue-specific variations in methylated transcripts ^23^. Our data also provide further evidence of m^6^A as a crucial regulator of synaptic organization and function in the adult brain.

### The m^6^A landscape in the adult human brain reveals a conserved enrichment of transcripts linked to synaptic function

Next, we decided to profile the m^6^A distribution across transcripts of the human brain employing postmortem tissue of the cingulate cortex (CC) from 5 non-demented individuals. In the human CC we found that in 22.8% of all expressed transcripts (3625) at least one m^6^A peak could be detected (Figure 2A). This corresponded to 11672 m^6^A detected peaks, with an average of 3.17 peaks per methylated transcript (Figure S3A). Similar to what is observed in mice, m^6^A peaks fell predominantly along the CDS and 3’UTR with a marked peak in the vicinity of the stop codon (Figure 2A, B). The consensus motif sequence for m^6^A marks DRACH, was consistently found enriched in the detected methylation peaks, confirming the specificity of the meRIP-seq method (Figure 2C). Human methylated transcripts belong to various molecular pathways, among them gene expression regulation, RNA metabolism, neural development and synaptic function (Figure S3B).

**Figure 2.**
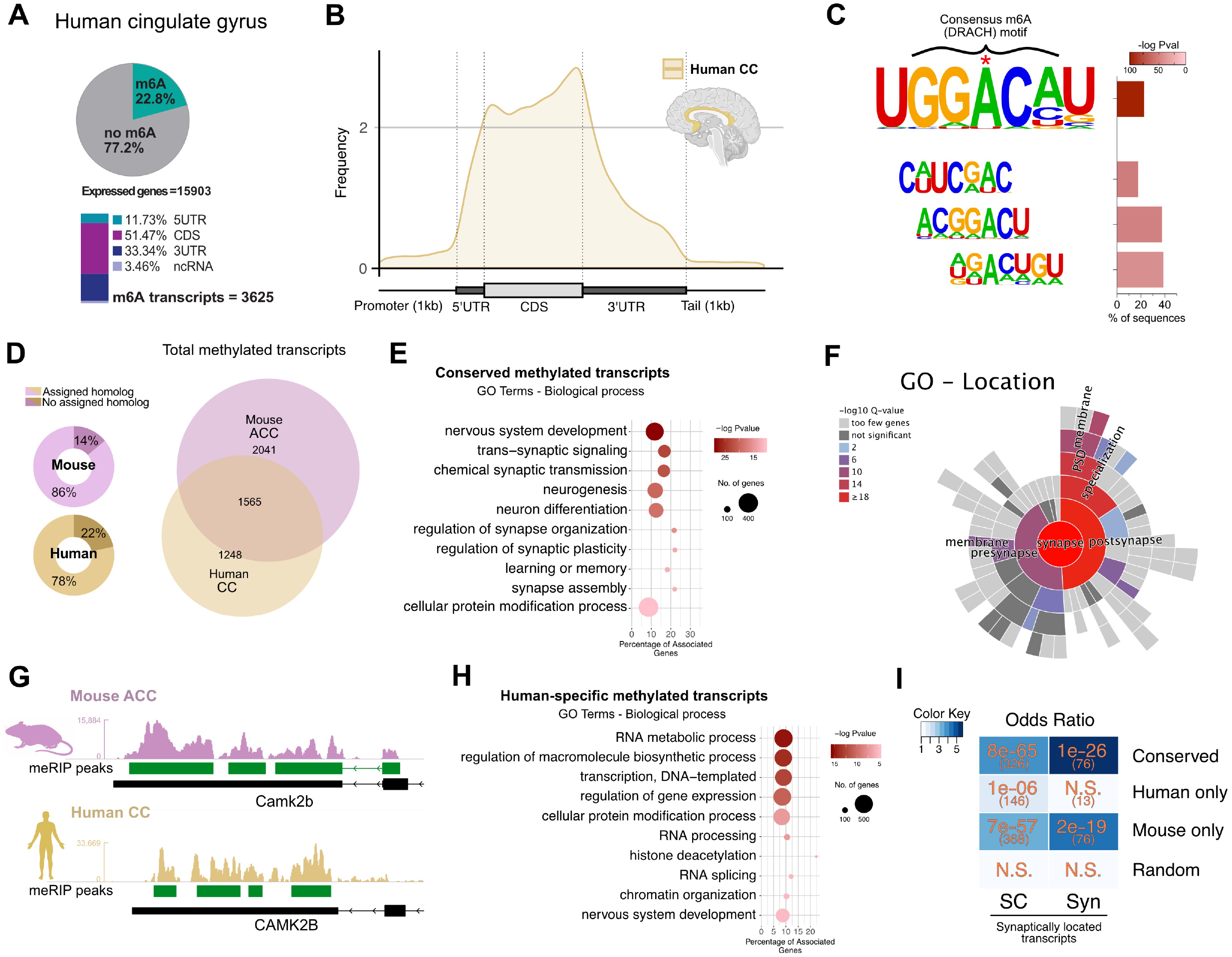
Conserved m^6^A marks between mouse and human. **A**. Percentage of methylated transcripts in the human CC, calculated against the background expression in the corresponding inputs; peak location by annotated region, percentages calculated from total m^6^A peaks. **B**. Distribution of m^6^A peaks along mRNA features in the human CC. **C**. Enriched motifs detected in the m^6^A peaks, showing the consensus m^6^A DRACH motif, where D=A,T or G, R=A or G, and H=A,T or C. **D**. Mouse/human genes with known homologs in human/mouse, respectively, were used to compare methylated transcripts across species. Overlap of methylated transcripts in the adult mouse ACC with respect to the human CC, from those genes with an assigned homolog. **E**. Enriched Gene Ontology categories, Biological process for genes methylated in both mouse ACC and human CC (conserved). **F**. Synapse-specific location GO term enrichment for conserved methylated transcripts. **G**. Representative coverage tracks showing conserved m^6^A sites along the 3’ end of homologous transcripts in the mouse ACC and human CC (Camk2b/CAMK2B). Tracks show coverage values for m^6^A-RIP normalized for the corresponding inputs and library size. Scale in RPM. **H**. Enriched GO categories, Biological process for genes methylated only in the human CC. **I**. Odds ratio showing the association between overlapping transcripts in conserved, human- and mouse-specific transcripts, compared to synaptically located RNAs, as published by Epple, et al. Color scale represents the numerical value of enrichment (odds ratio), numbers in orange correspond to the p value for the corresponding overlap, numbers in brackets show the size of the overlap. N.S.= not significant (p value ≥ 0.05). SC= RNAs detected in the synaptic compartments of microfluidic chambers; Syn= RNAs detected in synaptosomes. Random corresponds to 2000 randomly selected brain-expressed human genes. ACC - anterior cingulate cortex, CC - cingulate cortex

To compare methylated transcripts between brain regions across species in mouse and human we used the dataset previously generated for the mouse ACC and compared it with the human CC, both representing cortical brain regions. All transcripts with an assigned homolog in the corresponding species in NCBI’s Homologene database were used in this comparison, which accounted for the vast majority of all m^6^A transcripts (86% in mouse and 78% in human, Figure 2D). More than half (55%) of all methylated transcripts in human with an assigned homolog had their corresponding transcript in mouse methylated too (Figure 2D, Suppl. Table 5).

Functionally conserved transcripts were very strongly enriched for GO pathways linked to synaptic plasticity such as trans-synaptic signaling, regulation of synapse organization or learning and memory (Figure 2E). Furthermore, SYNGO analysis revealed that such transcripts are enrichment for synaptic location (Figure 2F) and function (Figure S3C).

Interestingly, not only the transcripts themselves were conserved in their methylation status, but also the location of methylation marks was in many cases conserved too. In the majority of them, annotated m^6^A peaks fell within the same region of the corresponding homologous human/mouse transcript counterpart (Figure 2G, S3D).

In addition to these conserved methylated transcripts, both the human CC and the mouse ACC had a subset of transcripts uniquely methylated in a species-specific manner. In the case of the mouse ACC these transcripts corresponded to genes involved in neurogenesis, the regulation of signal transmission and synaptic function, albeit with considerably less significant enrichment as in conserved transcripts (Figure S3E). Strikingly, transcripts uniquely methylated in the human CC showed a remarkable enrichment for genes associated with the regulation of gene expression, chromatin organization and RNA metabolism (Figure 2H), furthermore, they showed no synaptic localization (Figure 2I, Figure S3F). Supporting this functional specificity of conserved and non-conserved methylated transcripts, synaptically located mRNAs – both detected in synaptic compartments in microfluidic chambers and in synaptosomes – are consistently significantly overrepresented in the population of conserved and mouse-specific methylated transcripts. In contrast, human-specific methylated transcripts show low or no enrichment for synaptically located mRNAs, comparable to what could be expected by chance (Figure 2I).

These results indicate that the regulation of synaptic organization, function and plasticity through m^6^A marks is a conserved mechanism in the adult mammalian cortex. Moreover, species-specific differences in the methylation status of certain transcripts suggest that m^6^A marks are an evolutionarily dynamic regulatory mechanism, with certain populations of transcripts undergoing tissue- and species-specific labeling.

### m^6^A changes in models for cognitive decline and human AD patients

Our data supports the view that m^6^A-mediated regulation of synaptic function and plasticity is a key mechanism in the maintenance of homeostasis in the adult mammalian brain. To further explore this, we decided to study the m^6^A landscape during cognitive decline and choose age-associated memory impairment in mice as model system. Previous studies have reported that age-associated memory can be observed already in 16-months old mice, while at this stage only minor changes in neuronal gene-expression are detected ^39 40 41 42^. We reasoned that the comparison of 3 vs. 16 months old mice would thus allow us to test if changes in m^6^A RNA methylation may precede changes in gene-expression, as it had been reported for example for heart failure, which similar to the brain represents a disease affecting an excitable and mainly post-mitotic tissue ^43^. To this end we collected the brain subregions (ACC, CA1, CA3, DG) from 3 (young) and 16 (old) months old mice and performed meRIP-seq analysis (Figure 3A). In line with previous observations from bulk hippocampus, a differential expression analysis between old and young samples in the aforementioned subregions revealed comparatively mild changes (FC > 1.2, FDR≤0.05), ranging from 40 differentially expressed genes (DEGs) in the DG to 115 in the ACC (Figure 3B, Suppl. Table 6). DEGs were not significantly enriched for specific GO categories and no specific pathway was considerably affected (Figure S4A). In contrast to the transcriptome, the epi-transcriptome of these same tissues showed remarkable changes, when m^6^A levels in old mice were compared with the corresponding subregions in the young brain. Using the same cutoffs as for differential expression, a much larger number of genes showed differences in the methylation levels of at least one m^6^A peak along their transcript, consistently across all replicates (Figure 3B). The DG showed the most widespread changes, with 1971 transcripts differentially methylated, followed by the CA1 with 1557, ACC with 1373 and CA3 with 743 (Suppl. Table 7).

**Figure 3.**
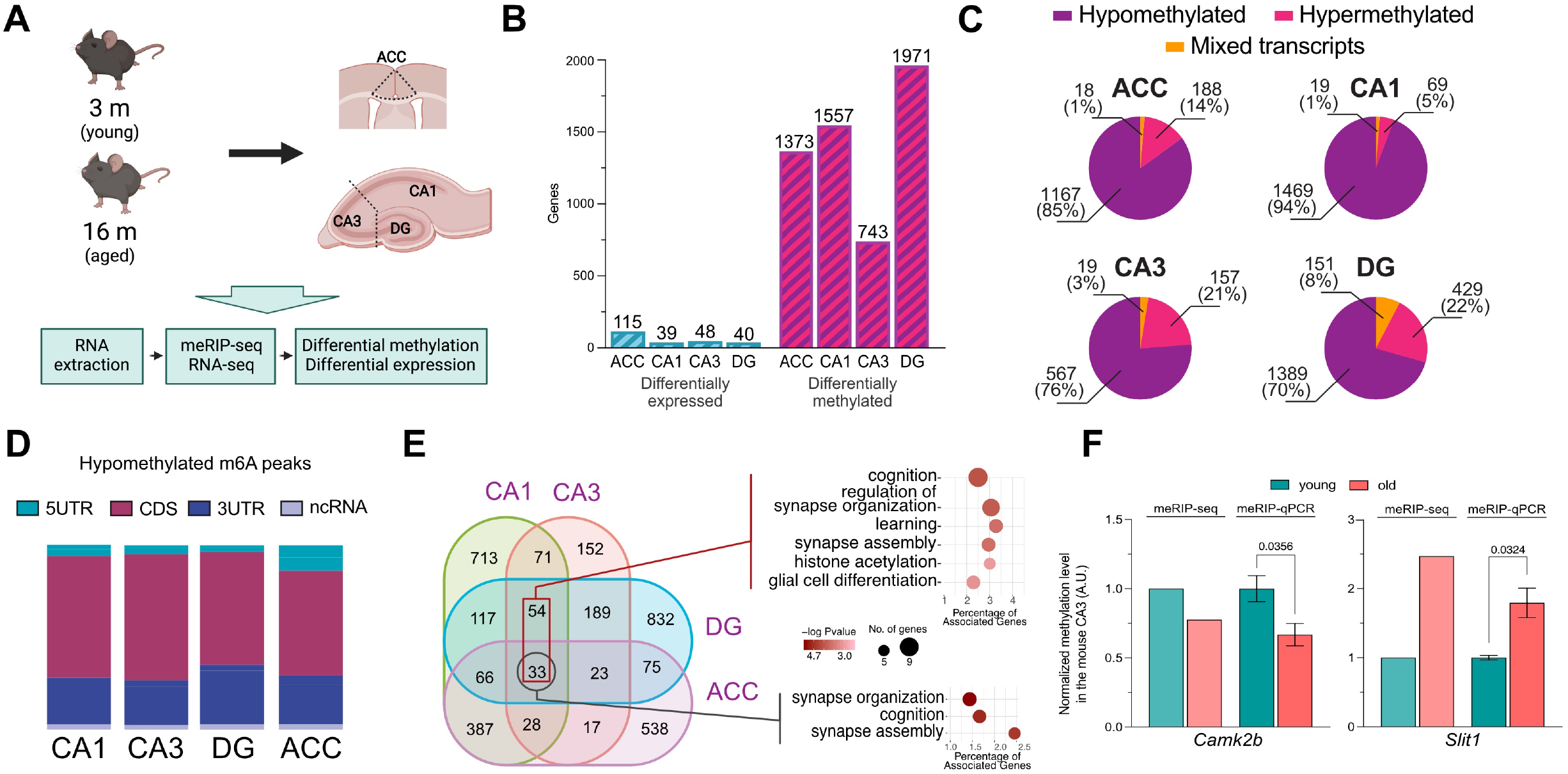
Tissue-specific m^6^A changes in the aging mouse brain. **A**. Workflow for differential methylation analysis in aging mouse brain. **B**. Bar graph showing the number of differentially expressed and differentially methylated genes detected in the corresponding brain subregion, applying equal cutoffs for fold change and adjusted p value (FC > 1.2, padj ≤ 0.05). **C**. Proportion of methylated transcripts containing peaks with only reduced methylation levels in aging (hypomethylated), only increased methylation (hypermethylated) or a mixture of decreased and increased (mixed) peaks in all brain subregions. **D**. Annotated distribution of significantly hypomethylated peaks across transcripts for all brain subregions. **E**. Overlap of hypomethylated transcripts across hippocampal subregions, highlighting the unique transcripts as well as common transcripts in the hippocampus. GO terms biological process for the 87 common transcripts across the hippocampus. **F**. qPCR validation of two differentially methylated genes. The graph shows the FC in methylation in the probed region as detected by meRIP-Seq and meRIP-qPCR. Columns show the mean +/-SEM of 4 independent replicates per condition. Statistical significance was determined by Student’s t test and the p value is displayed above the comparison. ACC. ACC - anterior cingulate cortex, DG - dentate gyrus, 5UTR - 5’ untranslated region, 3UTR - 3’ untranslated region, CDS - coding sequence, ncRNA - non-coding RNA

A total of 1698 peaks were detected as differentially methylated in the ACC, 2136 in the CA1, 811 in the CA3, and 2468 in the DG (Figure S4B). In all subregions, differentially methylated transcripts averaged 1.24 peaks per transcript (Figure S4C). At this level, sites of decreased m^6^A (hypomethylated) greatly outnumbered those with increased m^6^A (hypermethylated) across all brain subregions with the most striking changes occurring in the ACC and CA1 (Figure 3B). While the vast majority of differentially methylated transcripts (92-99%) showed consistent changes in methylation, in a few cases transcripts contained increased as well decreased m^6^A peaks (mixed transcripts, Figure 3C). In all subregions, the bulk of differentially methylated transcripts showed consistent hypomethylation, with up to 94% of the total in the CA1 and 85% in the ACC belonging to this group. In contrast, only a small fraction of transcripts was consistently hypermethylated, with this population being most numerous in the CA3 and DG, with 21% and 22%, respectively (Figure 3C). Some variability was also observed in the magnitude of change across brain subregions, with the CA1 and ACC displaying a more dramatic reduction in m^6^A, compared to the CA3 and DG (Figure S5A).

The location of m^6^A marks along the transcript has been associated with distinct fates for the labeled mRNAs. To determine whether aging-associated changes were favoring certain regions of labeled transcript, differentially methylated peaks were annotated according to their location. Like it is the case with the baseline methylated regions, differentially methylated peaks are enriched along the gene body, stop codon and 3’UTR (Fig.1B, C, Fig. 3D). Interestingly, the ACC shows slight enrichment of differentially methylated peaks within the 5’UTR while hypermethylated peaks were slightly enriched in the vicinity of the stop codon, when compared to the other analyzed subregions (Fig. 3D, Figure S5B). In addition, hypomethylated peaks in the DG displayed a slightly stronger increase in their location surrounding the stop codon as well. However, hypermethylated peaks showed a considerably larger variability in their location within mRNAs, in large part due to their smaller numbers (Figure S5B-C).

When comparing the hippocampus, 87 mRNAs are detected as hypomethylated in all hippocampal subregions, whereas for shared hypermethylated transcripts the number is 2 (Figure 3E, Figure S5D). Despite this limited commonality, the shared hypomethylated transcripts are highly enriched for pathways associated with cognition, learning and synaptic organization, showing that despite high tissue specificity, hypomethylation affects certain common pathways in the hippocampus (Figure 3E).

Similarly, when comparing hypomethylated transcripts amongst hippocampal subregions and the ACC, only 33 of them could be detected in all tissues. The largest group of commonly hypomethylated transcripts is found in the ACC and CA1, where 387 of them are hypomethylated in both tissues (Figure S3F, Suppl. Table 8). Despite the limited commonality of individual transcripts hypomethylated across all regions, the pathways affected by decreasing methylation showed a remarkable similarity in the hippocampus and ACC (Figure 3E; Figure S5E). Significantly differentially methylated regions detected at the sequencing level could be validated independently by meRIP-qPCR (Figure 3F).

Taken together, these data identify massive changes in m^6^A levels in the aging brain at a time point when first memory impairment is observed and gene-expression changes are comparatively moderate. The vast majority of the affected transcripts exhibit m^6^A hypomethylation and represent genes linked to synaptic plasticity.

These data hint towards a role of this mark in the development of age-associated cognitive decline, a role that would be in line with reported observations of the involvement of m^6^A in learning and memory. Considering the commonality in m^6^A transcripts in the brains of mice and humans (see Figure 2) we decided to study whether m^6^A changes could be associated with cognitive impairment in humans. Thus, we analyzed the m^6^A-epitranscriptome during Alzheimer’s disease (AD), the most common cause of dementia in the elderly. Postmortem human cortex samples from AD patients were matched with corresponding non-demented controls (NDC) and analyzed by meRIP-seq. At the gene expression level, we detected a total of 185 genes as differentially expressed, with 100 of them being upregulated and 85 downregulated (FC > 1.2, FDR ≤ 0.05, Figure 4A, Figure S6A). GO terms for upregulated genes show an enrichment of regulators of the Wnt signaling pathway, whereas downregulated genes were not associated with a given pathway. It is worth noting that no genes associated with the m^6^A machinery were amongst those with significant expression changes in this dataset (Figure S6A).

**Figure 4.**
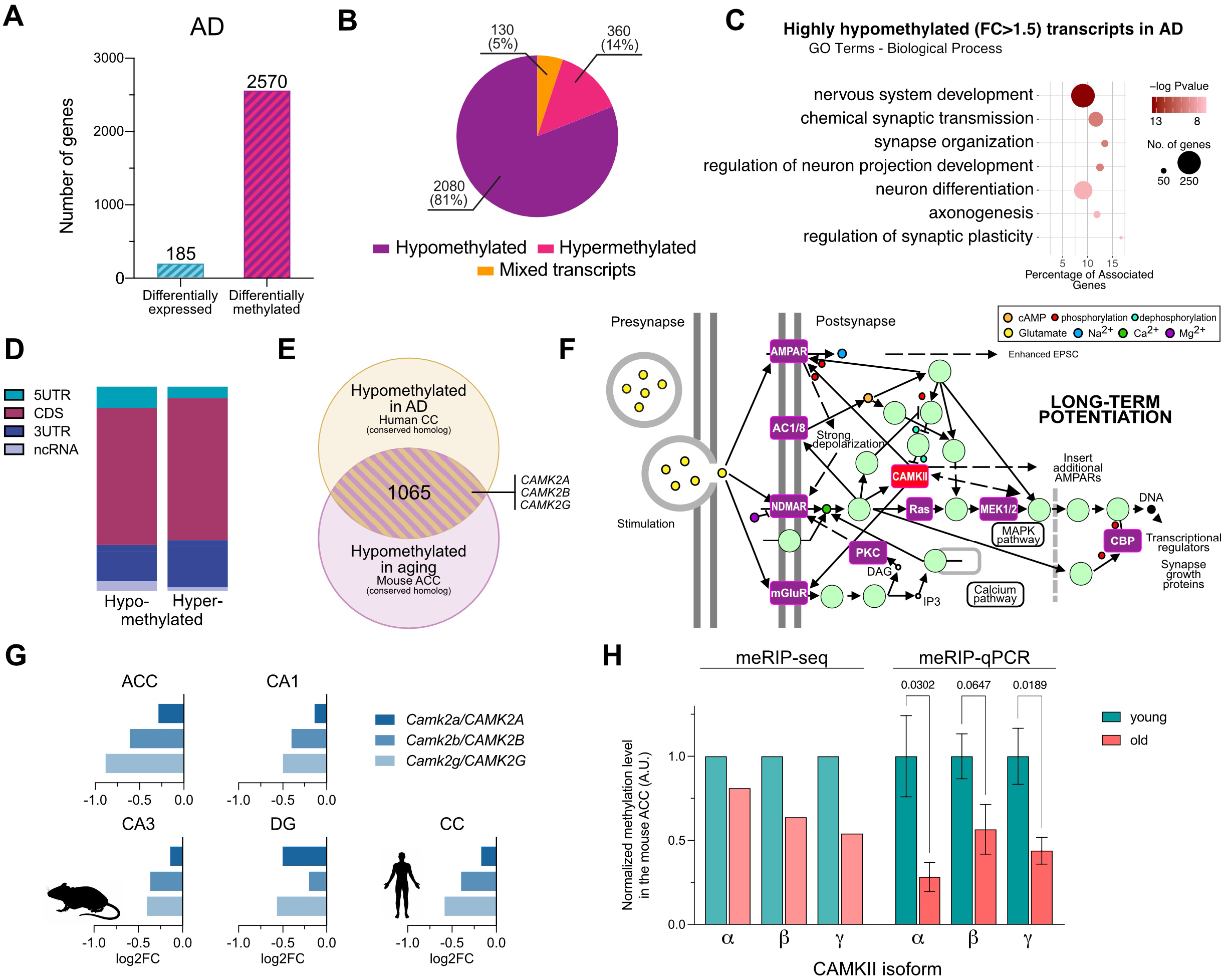
Epi-transcriptome changes in neurodegeneration and aging. **A**. Comparison of differentially expressed and differentially methylated genes in AD vs control samples applying equal cutoffs for fold change and adjusted p value (FC > 1.2, padj ≤ 0.05). **B**. Proportion of hypomethylated, hypermethylated and mixed transcripts in AD samples. **C**. Enriched GO categories Biological process for hypomethylated transcripts (FC > 1.5) in AD. **D**. Annotated distribution of differentially methylated peaks in AD along transcripts. **E**. Overlap of all significantly hypomethylated peaks in the aged mouse ACC and AD human CC, highlighted are CaMKII isoforms. **F**. KEGG pathway *Long-term potentiation* (hsa04720), highlighted in purple are pathway components that are commonly hypomethylated in aging and AD, in red is CAMKII. **G**. Changes in methylation in individual isoforms of CaMKII in the aged mouse brain and in the human CC in AD. Each bar represents the methylation site in the 3UTR of the corresponding transcript closest to the stop codon. **H**. qPCR validation of the hypomethylated region in the 3UTR of the displayed CAMKII isoforms. The graph shows the FC in methylation in the probed region as detected by meRIP-Seq and meRIP-qPCR. Bar shows the mean +/-SEM of 6/4 (young/old) independent replicates, p value is displayed above the comparison. Statistical significance was evaluated by Student’s t test with Welch’s correction for unequal variances. AD - Alzheimer’s disease, ACC - anterior cingulate cortex, DG - dentate gyrus, 5UTR - 5’ untranslated region, 3UTR - 3’ untranslated region, CDS - coding sequence, ncRNA - non-coding RNA.

In stark contrast to the changes at the gene expression level, the differential methylation analysis of meRIP-seq shows massive changes in m^6^A levels in AD. More than 2500 genes are detected as differentially methylated using the same FC and FDR cutoffs as the differential expression analysis (Figure 4A, Suppl. Table 9). These correspond to 3288 differentially methylated peaks (2568 hypo- and 424 hypermethylated) with an average of 1.26 m^6^A peaks per differentially methylated transcript (Figure S6B). Of them, the vast majority are exclusively hypomethylated transcripts (81%) with a smaller fraction (14%) showing only hypermethylation and the remaining 5% having both hypo- and hypermethylated regions (mixed transcripts, Figure 4B). Hypomethylated transcripts showed a very strong enrichment of genes associated with developmental processes, neuron projection and the regulation of synaptic transmission and plasticity, GO categories that also were highly affected by m^6^A changes during aging in the mouse brain (Figure 4C). The location of these differentially methylated marks did not favor any specific region of the affected transcripts (Figure 4D. Figure S6C).

These data are similar to the observed changes of m^6^A levels in the aging mouse brain. In fact, there was a considerable overlap between the populations of hypomethylated transcripts in the aged mouse brain and the human AD brain and more than 1000 hypomethylated m^6^A transcripts were detected in both species (Figure 4E, Suppl. Table 10). Among these transcripts, the majority has well described roles in the regulation of synaptic function, learning and plasticity (Figure S6E). Furthermore, there was a very significant overlap between these transcripts and RNAs previously described as synaptically located, as well as known synaptic methylated transcripts (Figure S7) ^38 27^. In this group, numerous components of pathways associated with the regulation of plasticity - like long-term potentiation (LTP) - and disease are highly overrepresented (Figure 4F, Figure S6F). Within this subset we found multiple isoforms of one of the best described subfamilies of synaptic plasticity-associated proteins, the Calcium/calmodulin-dependent protein kinase type II (CamkII), namely *Camk2a, Camk2b* and *Camk2g* (Figure 4E-G, Figure S6F). CaMKII, and especially the α and β isoforms, are central for memory formation and learning ^44^. The corresponding transcripts were characterized by a consistent hypomethylation in the aging mouse and human AD brain (Figure 4G, Suppl. Table 11) a finding that was confirmed by qPCR (Figure 4H).

### Decreased m^6^A levels affect the local synthesis of the plasticity-related protein CaMKII

Next, we wanted to further elucidate the functional consequences of reduced m^6^A methylation. The fact that we see comparatively few changes in gene-expression while substantial m^6^A-hypomethylation is observed in aged mice or AD brains, suggest that the m^6^A changes detected in our experimental settings may not impact on transcript stability, a process that has been linked to m^6^A RNA-methylation ^45^. In line with this hypothesis there was no obvious correlation between m^6^A and transcript changes in any of the analyzed tissues (Figure S8A-E). This finding was further corroborated by our observation that levels of histone 3 tri-methylation at lysine 3 (H3K36me3), a repressive histone-mark that had been linked to transcription-dependent changes in N^6^A RNA-methylation ^46^, were similar when comparing hippocampal tissues samples from young and old mice via ChIP-sequencing (Figure S8F,G). M^6^A labels on mRNA are also known to play a role in regulating the transport of certain synaptically-located transcripts, as well as on the somatic translation of plasticity-related genes ^27 26 16^. To determine whether these mechanisms could be acting downstream of the changes in m^6^A levels during aging and AD, we first isolated synaptosomal compartments from the hippocampi of young and old mice and performed RNA-seq on the resulting synaptic mRNA population (Figure S9A). Similar to the analysis of bulk tissue (See Fig. 4), we detected comparatively few differentially expressed transcripts in synaptosomes when analyzing the data from 3 vs 16-months old mice (3 transcripts up-regulated and none down-regulated) (Figure S9B). Also, neither for these nor for any of the detected transcripts was there a correlation to their methylation status (Figure S9C). In sum, these data suggest that aberrant transport of transcripts from the soma to the synapses may not be the major consequence of m^6^A hypomethylation in our disease models. Another process linked to m^6^A RNA-methylation is mRNA translation ^15^. Thus, we performed polysome sequencing on young and old hippocampal tissue samples. Differential binding analysis identified 83 genes to be differentially translated during aging (Figure S9D, E) but there was no significant overlap to the transcripts affected by differential m^6^A methylation (Figure S9F).

Since m^6^A RNA-methylation has been associated with local protein synthesis (LPS), the analysis of bulk tissue via polysome-seq might not be sensitive enough to detect the respective changes. In fact, m^6^A was shown to control axonal protein synthesis in motorneurons ^47 26 48^. Thus, we hypothesized that the observed changes in m6A levels may affect synaptic protein synthesis. To address this experimentally, we opted to use a primary neuron model to evaluate the effect of reduced m6A levels on LPS at the synapse. Since there are no suitable high-throughput methods to assay synaptic LPS, we decided to evaluate its rate and location by studying the synthesis of CAMKII via a puromycin-based proximity ligation assay (puro-PLA). We chose CAMKII since it is a well-described synaptically-located transcript that is known to be locally-translated, plays a key role in memory function and was hypomethylated in the aging mouse and human AD brain ^47 26 48 49 38^.

To reduce m^6^A levels, the methyltransferase Mettl3 was knocked down (KD) on primary hippocampal mouse neurons. The KD of Mettl3 via siRNA has been reported to be challenging in primary neurons ^27^. Indeed we observed only partial effects despite high concentration of siRNA probes (Figure S10A,B). To improve on this, we decided to employ another technology, namely LNA GAPmers, at lower doses and for longer treatment periods. Primary neurons transfected with an LNA GAPmer targeting Mettl3 packaged in lipid nanoparticles (LNPs) at day in vitro (DIV) 7 showed an almost complete reduction in the Mettl3 mRNA level (> 95%) when measured 3 days later (Figure 5A, Figure S10C). However, further 3 days of culture were necessary to sufficiently decrease the METTL3 protein and m^6^A levels (Figure 5B-D). Having established successful reduction of m^6^A levels we studied PLA of CAMKII via the puro-PLA assay. Puro-PLA depends on the use of the antibiotic puromycin for the labeling of nascent protein chains and N-terminal primary antibodies to detect sites of translation through proximity ligation (Figure 5E, Figure S10D) ^50^. A cycloheximide pretreatment was also applied to improve the spatial localization of sites of protein synthesis ^51^. Puromycin labeling and translational arrest were confirmed in treated neurons (Figure S10E). DIV 13 primary neurons that had been treated at DIV 7 with either a Mettl3 KD or control GAPmer were processed for puro-PLA using an antibody that detects the N-terminal of CAMKII α, β and γ (Figure 5E,F). Puro-PLA-treated neurons were imaged by confocal microscope and the PLA punctae automatically detected and quantified, the synaptic marker Synaptophysin (SYP) was used to determine the synaptic localization of detected punctae (Figure 5F). Neurons with reduced levels of m^6^A (Mettl3 KD) showed a reduction of PLA punctae in dendritic projections (Figure 5F, Figure S11A). Quantitative analysis revealed that the total number of PLA punctae in the whole neuron was not significantly reduced (Figure 5H, Figure S11B). The number of detected synapses was also not significantly changed in response to decreased m^6^A levels (Figure S11C,D). However, when looking at the proportion of CaMKII-PLA punctae detected in vicinity to SYP+ synaptic compartments, the Mettl3 KD-treated neurons showed significantly decreased numbers (Figure 5F,H). To rule out the possibility of these differences are a consequence of decreased mRNA transport to synaptic compartments, we used a previously established custom-made microfluidic chamber culture system to isolate synaptically located transcripts (Figure S2E) ^38^. Mettl3 KD treatment on the somas of the cultured neurons showed no significant effect on the amount of mRNA located in synaptic compartments for *Camk2a, Camk2b* and *Camk2g* (Figure 5I, Figure S11E).

**Figure 5.**
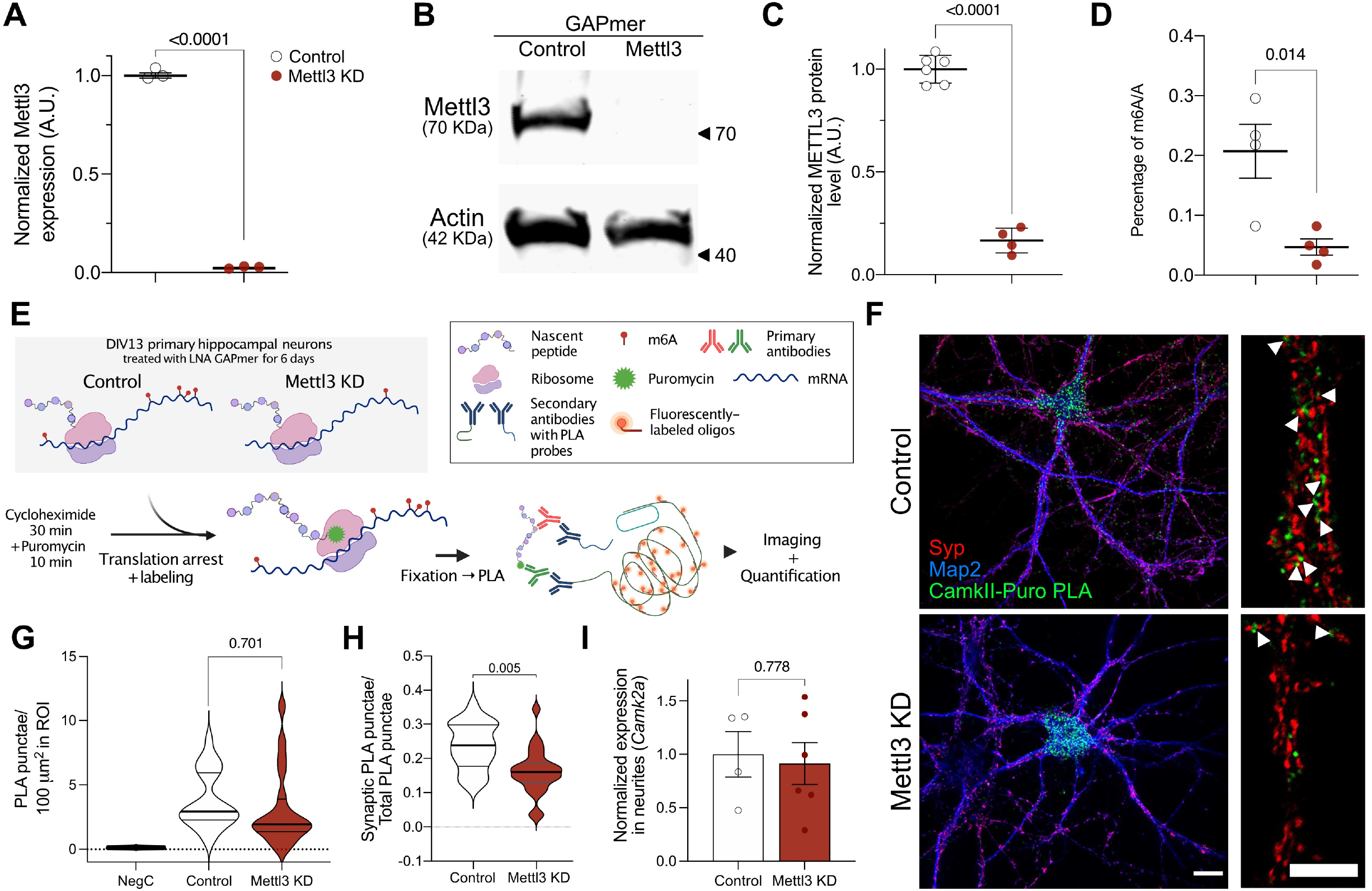
m^6^A changes influence the synaptic protein synthesis of CaMKII. **A**. Validation of GAPmer-mediated knockdown of *Mettl3* at the transcript level via qPCR **B. METTL3** protein level in response to GAPmer-mediated knockdown of *Mettl3* were analyzed via Western blot. **C**. Quantification of (B). **D**. Analysis of global m^6^A levels upon GAPmer-mediated knockdown of *Mettl3.* Graphs in A, C and D display the mean +/-SEM of each condition. Each data point represents one independent replicate, statistical significance was determined by Student’s t test. **E**. Schematic illustration of the Puro-PLA labeling used to quantify the synthesis of CaMKII in primary neurons. **F**. Representative images of Control and Mettl3 KD primary hippocampal neurons displaying the localization of CaMKII-PLA punctae Scale bar = 20μm. CaMKII-PLA signal in green, Synaptophysin in red, Map2 in blue. Zoomed-in images show a large magnification image of a representative dendrite (Scale bar = 10μm). Arrowheads show sites of CAMKII LPS in the close vicinity of synapses. **G**. Total number of detected PLA punctae in treated neurons. Negative control (NegC) was not treated with puromycin before being processed for PLA. **H**. Synaptically-located CaMKII-PLA punctae in control and Mettl3 KD-treated neurons. Graphs in G and H show the mean of 3 independent experiments, for each experiment 7-13 neurons were imaged and analyzed, individual data points were used to generate the violin plot. Quartiles are marked by gray lines. **I**. Normalized *Camk2a* mRNA levels in the synaptic compartments of treated and control primary cultures in microfluidic chambers. Dots in I represent individual independent replicate cultures. Statistical significance determined by Student’s t test. For all panels, p values are displayed above the corresponding comparisons.

These results support the idea of that m^6^A changes in aging and neurodegeneration affect synaptic function and memory through the regulation of the synaptic LPS of plasticity-related mRNAs.

## Discussion

In this study, we aimed to further disentangle the role of m^6^A function in the adult brain and analyzed the m^6^A epitranscriptome in mice and humans. We found that 40-44% of the detected transcripts in the different hippocampal subregions in mice carried m^6^A marks. These data are in line with recent studies in which different brain regions or bulk hippocampal tissue had been analyzed for m^6^A RNA methylation ^52,53 36^. m^6^A-labeled mRNAs in the adult hippocampus were strongly enriched for genes associated with the regulation of synaptic function and plasticity. This enrichment holds true for all hippocampal subregions, with a very considerable overlap between the populations of labeled transcripts in the CA1, CA3 and DG subregions. We also compared the methylated transcripts between the ACC and the hippocampus, two more distantly related brain regions at the structural and functional level, and observed that 61% of the m^6^A transcript could be detected in both brain regions. These results go in accordance with previously published data that reported a considerable amount of tissue specificity in the populations of transcripts that are labeled by m^6^A ^23 54 53^. While noticeable difference in the m^6^A landscape were detected between brain subregions, the location of m^6^A marks were mostly identical within all hippocampal regions and the ACC. The commonly N^6^A methylated transcripts were mainly involved in synaptic signaling and structure, suggesting the general importance of certain m^6^A-dependent regulatory networks in synaptic plasticity. Moreover, the m^6^A transcripts commonly detected across brain regions exhibited a large over-representation of mRNAs that are localized to synapses, which is in line with data suggesting that m^6^A transcripts are specifically enriched in synaptosomes ^27^. The m^6^A transcripts specific to the different hippocampal subregions were linked to more general cellular processes such as the negative regulation of protein complex assembly in the CA1 region, processes related to development and RNA processing in the DG and for example protein transport in the CA3 region. When comparing the m^6^A landscape from the mouse anterior cingulate and human cingulate cortex, we observed that 56% of the methylated transcripts found in humans, were also detected in mice. This is remarkable, when considering that a similar degree of conservation is observed when the anterior cingulate cortex is compared to the hippocampus within the same species (61% in mice). These data are in agreement with a previous study that found 62% overlap between m^6^A transcripts of the mouse and human cerebellum ^53^. The commonly methylated transcripts detected in the mouse and human cortex were linked to the regulation of synaptic function and plasticity and showed a strong overrepresentation of transcripts found at synapses. Interestingly, the m^6^A transcripts specific to mice were enriched for GO-terms related to synaptic plasticity, while the methylated transcripts specific to the human cingulate cortex were also strongly enriched for GO-terms linked to RNA processing and gene-expression control. These data may suggest that the orchestration of synaptic plasticity is an evolutionary conserved mechanism in mammals, while a role of m^6^A RNA-methylation in gene-expression control, gained importance in the human brain. However, care has to be taken when comparing data from cortical regions in mice and humans.

To study the epi-transcriptome in the context of cognitive dysfunction we employed aged mice and human AD patients. During aging, the onset of mild cognitive impairment represents a hallmark of the transition between normal aging and pathology ^42 55 56^ and previous data demonstrated that a significant memory impairment can first be detected when comparing mice at 3 vs. 16 months of age ^57 42^. In addition to the fact that this animal model does not dependent on the expression of a transgene, previous studies showed that only minor changes in gene-expression are observed in the hippocampus and cortex when comparing 3 to 16-month-old mice ^39^, making these animals a suitable model to study changes in m^6^A transcripts. We observed a massive hypomethylation across multiple transcripts in all investigated brain regions, while comparatively mild change in gene-expression were observed. This in agreement with data from other postmitotic tissues, namely the heart, where during the pathogenesis of heart failure massive changes in m^6^A hypomethylation precede change in gene-expression ^43^. Amongst the different brain regions affected by aging, there was a noticeable tissue specificity, as many region-specific changes could be detected. These results show that different brain regions undergo distinct changes during aging, but the overall pathways affected by m^6^A changes remain similar and were linked to GO-pathways such as cognition and synapse organization. These data suggest that loss of m^6^A RNA-methylation is an early event in the aging brain that coincides with the onset of memory impairment. This view is supported by previous data showing that a knock-down of the m^6^A demethylase FTO in the prefrontal cortex of mice results in an improved consolidation of fear memories ^58^. Similarly, loss of the m^6^A reader YTHDF1 which has been linked to enhanced translation, leads to impairment of hippocampal LTP and memory formation in mice ^16^, another recent study analyzed m6A levels in the brains of 2 week, 1, 1.5, 6.5 and 13 months-old mice. The authors observe comparatively milder changes than in our study and the affected transcripts were mainly characterized by altered m6A levels within the UTR that increased from 1.5 to 13 month of age ^36^. Since animals at 13 months of age do not exhibit detectable memory impairment ^59^, these changes may represent compensatory mechanisms. In line with this interpretation, the affected pathways were linked for example to cellular stress signaling. The same study also analyzed m^6^A levels in the brains of 6 months old 5xFAD mice, a mouse models for amyloid deposition as it is observed in AD ^36^. Here, decreased m^6^A levels were observed when comparing wild type to 5xFAD mice and the affected transcripts were linked to GO-terms such as synaptic transmission. On the basis of previous data showing that 5xFAD mice display memory impairment at 6 months of age ^60^ these data are in agreement with our observations. Nevertheless, more research is needed to elucidate the dynamics of m^6^A marks across the transcriptome of the aging and diseased brain.

Additional support for the hypothesis that cognitive decline is accompanied by m^6^A hypomethylation of transcripts important for synaptic function stems from our analysis of postmortem human brain samples from AD patients. The AD brains showed significant changes in methylation, with the majority of transcripts being hypomethylated. When compared to the hypomethylated peaks in the aging mouse ACC, striking similarities could be found for the affected transcripts and in the corresponding pathways. Among them, the regulation of synaptic plasticity, particularly long-term potentiation, as well as multiple neurodegeneration-associated pathways were strongly affected.

The finding that AD is associated with m^6^A hypomethylation is in agreement with a recent study in which a strong decrease in the levels of the main m^6^A methyltransferase METTL3 was observed in the hippocampus of AD patients at the mRNA and protein level ^30^. Reduced expression of m^6^A writers could indeed be one mechanism to explain lower m^6^A levels in AD. In line with this view, knock-down of *Mettl3* exacerbated Tau pathology in a Drosophila model for AD ^36^. It should be mentioned that the role of N^6^A RNA-methylation in neurodegenerative disease may be more complex. For example, a recent study observed increased m^6^A levels in a mouse model for Tau-pathology and in the brains of human AD patients ^32^. However, these data are based on a semi-quantitative analysis m^6^A immunostaining within the soma, which is difficult to compare to sequencing-based approaches. Similarly, another recent study reported an increase of bulk m^6^A and METTL3 levels, while FTO protein levels were decreased in the hippocampus and cortex of 9 months old APP/PS1 mice ^34^. The fact that the analysis of sequencing-based vs. bulk analysis of m^6^A levels currently appear contradictive, may indicate that there is an RNA-species which undergoes hypermethylation in neurodegenerative diseases, that is not captured by the current sequencing approaches but dominates the analysis of bulk m^6^A levels. For example, recent evidence hints to an important role of N^6^A-methylation of pre- and mature microRNAs ^61 62^ and it will be interesting to study microRNA methylation in brain diseases. In addition, it will be important to study m^6^A levels in neuronal subcompartments. In fact, our data consistently show that m^6^A hypo-methylation occurs often within synaptically located plasticity-associated transcripts, pointing to a role of N^6^A RNA-methylation in the local translation of synaptic transcripts, a well-known phenomenon that ensures the supply of key proteins necessary for synaptic function and plasticity in response to stimuli ^63 64 65^. The function of m^6^A as a regulator of LPS has been shown in the axons of motoneurons ^47^ and since then has been theorized in other contexts but so far, no direct experimental evidence has been put forward to prove this link.

Our results show that a reduction in m^6^A caused by a decrease in Mettl3 expression, akin to the reductions observed during aging and AD, significantly impact the rate of protein synthesis of the plasticity regulator CAMKII in the vicinity of synaptic compartments, away from the soma.

Taken into account that we found *CamkII* transcripts to undergo m^6^A hypomethylation in both aging and AD and that *CamKII* was also among the list of transcripts that underwent hypomethylation in the cortex of 5xFAD mice ^36^, these data suggest a m^6^A-depedent mechanisms that orchestrates synaptic proteins synthesis and contributes to impaired synaptic plasticity when de-regulated.

Future research is needed to elucidate the precise mechanism by which m^6^A levels control the synaptic translation of transcripts. In this context it is noteworthy that the demethylase FTO was shown to locate at the synapse and that its levels decrease during learning ^24^. At the same time the m^6^A reader YTHDF1 is also located in synaptic compartments and its protein levels significantly increase following fear conditioning in the hippocampus ^28^. YTHDF1 was shown to promote translation ^15 28^ and although these data do not stem from synapses, this might be one mechanism by which reduced m^6^A levels affect synaptic protein synthesis. Furthermore, the knockdowns of *Ythdf1* negatively affects spine formation, long-term potentiation, and learning in a hippocampus-dependent manner ^26 28^. More recently, the m^6^A reader YTHDF3 as well as the eraser ALKBH5 have also been linked to the regulation of m^6^A at the synapse, further increasing the possibilities for regulation in such compartments ^54^. Of course, other m6A reader may also play a role in regulating synaptic mRNA translation directly or indirectly, via processes like degradation, transport or phase separation ^31 32 27^.

In conclusion, our data provide an important resource to the field and further elucidate the function of m^6^A in regulating learning and memory in the healthy and diseases brain by showing that m^6^A controls synaptic LPS and its relationship with decreased methylation of synaptic genes during aging and neurodegeneration.

## Materials & Methods

### Animals

3 months old (young) and 16 months old (old) male mice were purchased from Janvier Labs. Animals were housed under standard conditions. Experiments were performed according to the protocols approved by the Lower Saxony State Office for Consumer Protection and Food Safety (LAVES).

### Human AD tissue

A total of 12 post-mortem samples from the anterior cingulate cortex were obtained from the Netherlands Brain bank. The samples corresponded to 6 diagnosed AD patients (age 89.33 ±4.42 years, Braak and Braak stages IV, PMD 6:34 ±1:00) and 6 non-demented controls (age 86.33 ±3.25 years, Braak and Braak stages I-II, PMD 6:16 ±1:38), all individuals, except one AD patient were female. All experiments were approved by an ethics committee.

### RNA extraction

RNA used for sequencing was extracted from tissue using the NucleoSpin RNA/Protein Kit (Macherey-Nagel), according to the manufacturer’s instructions. RNA integrity was assayed by electropherogram in a Bioanalyzer using a total RNA Assay with a Pico/Nano Chip (Agilent).

### meRIP

RNA samples were processed as previously described for meRIP-seq ^43^. The rRNA-depleted RNA from four mice was pooled together for each replicate, for a total of three replicates per condition and fragmented for 8 minutes at 70°C to an average fragment size of ~80 nt using Fragmentation Reagents (Invitrogen). rRNA depletion, fragment size and RNA quality were controlled in the Bioanalyzer. 10 μg of anti-m^6^A antibody (Synaptic Systems) or rabbit IgG control (Millipore) were added to 50 μl of Protein A/G beads (Thermo) and incubated in 500 μl of IP buffer (0.2 M Tris-HCl pH 7.5, 0.5 M NaCl, Igepal 2%) for 4 hours at 4°C with constant rotation. 10 μg of fragmented RNA per replicate were used for the subsequent RIP, with 500 ng (5%) kept to serve as the input. The RNA was incubated with the antibody-beads conjugate, in 1 ml of IP buffer supplemented with 200 units SUPERase-in (Invitrogen) overnight (ON) at 4°C. Beads were washed 5 times with IP buffer and precipitated RNA was eluted with 6.7 mM m^6^A in 200 μl IP buffer for 1 hour at 4°C with agitation. Eluted RNA was cleaned before proceeding to library preparation.

### Library preparation and sequencing

Samples were prepared for sequencing using the TruSeq Stranded Total RNA Library Prep Kit (Illumina) or the SMARTer Stranded Total RNA Kit v2 - Pico Input Mammalian (Takara) according to the manufacturer’s instructions.

### Bioinformatic analysis of meRIP-Seq RNA-Seq

Raw reads were processed and demultiplexed using bcl2fastq (v2.20.2) and low-quality reads were filtered out with Cutadapt v1.11.0 ^66^. Filtered reads were mapped to the human (hg38) or mouse (mm10) genome using the STAR aligner v2.5.2b ^67^. The resulting bam files were sorted, indexed and the unmapped reads removed using SAMtools v1.9.0 ^68^. Methylation sites were determined using MeTPeak v1.0.0 ^69^ and differential methylation was assessed with Exomepeak v2.16.0 ^70^, an adjusted p value (padj, also termed FDR [False Discovery Rate]) cutoff of 0.05 and fold-change (FC) cutoffs of 1.2 or 1.5 were used as indicated in the text. For mouse samples, only consistently significantly differentially methylated peaks were used, unless indicated; for human samples, significantly differentially methylated peaks were used.

For RNA-Seq analyses, read counts were obtained with subread’s featurecounts v1.5.1 ^71^ from the bam files of input samples. Differential gene expression was determined by DESeq2 v3.5.12 ^72^ using normalized read counts and correcting for covariates detected by RUVseq v1.16.1 ^73^. Cutoffs of padj ≤ 0.05, FC ≥ 1.2 and BaseMean ≥ 50 were applied to the results.

For visualization, bam files of both IP and input samples were collapsed for PCR duplicates using SAMtools and IP samples were normalized to their corresponding inputs and to their library size using deeptools’ v3.2.1 ^74^ bamCompare. The resulting normalized tracks were visualized in the IGV Browser 2.9.2 ^75^.

### Gene ontology (GO) analyses

GO term enrichment analyses were performed using the App ClueGO v2.5.3 ^76^ in Cytoscape 3.7.2 ^77^, with GO Term Fusion enabled to collapse terms containing very similar gene lists. GO term tables for Biological process, Cellular component, Pathways and KEGG were produced and are labeled accordingly in the figures. Resulting enriched GO terms were visualized with a custom script using ggplot2 v3.3.5 ^78^, displaying the adjusted p value (padj) for the GO term, the number of genes from the list that belong to said term and the percentage of the total genes in the GO term that are present in the list. Synaptic GO enrichment analyses were performed with SynGO (v1.1, syngoportal.org)^37^.

### Additional bioinformatic packages and tools

Scripts and analysis pipelines were written in R (3.5.2)^79^. Peak annotation was performed with Homer v4.10.4 ^80^ and Annotatr v1.8.0 ^81^. Guitar plots were produced with the Guitar v1.20.1 ^82^ R package. Volcano plots were generated with plot.ly/orca v4.9.4.1 ^83^. Area-proportional Venn diagrams were produced with biovenn (www.biovenn.nl)^84^ and multiple lists comparisons performed with Intervene/UpSet (asntech.shinyapps.io/intervene/)^85^. Mouse/human homologs were determined by their annotation in NCBI’s Homologene database using the Homologene (v1.4.68.19.3.27) R package. Odds ratios and p values to determine significance in overlapped datasets were calculated with the GeneOverlap R package v1.18.0 ^86^. De novo motif analyses were performed with Homer’s findMotifsGenome and motifs containing the DRACH consensus sequence out of the top 10 most significant are displayed. KEGG pathway enrichment was produced with KEGG Mapper (www.genome.jp/kegg/mapper/)^87^. Microscopy images were preprocessed with Fiji ^88^ and quantification was automated in Cell Profiler (cellprofiler.org)^89^. Graphs, heatmaps and statistical analyses were performed on GraphPad Prism version 9.3.1 for Mac. Some custom figures were created with BioRender (biorender.com).

### qPCR

qPCR was performed as describe before^59^. Primer sequences are available in Supplementary Table 1.

### Hippocampal primary neuronal culture

Primary neurons were prepared as described recently^59^.

### siRNA/ LNA GAPmer transfection

Pre-designed control and Mettl3-targetting siRNAs were purchased from Origene, control and Mettl3-targeting LNA GAPmers were designed and purchased from Qiagen. 2 pmol of the corresponding control/Mettl3 siRNA/GAPmer were packaged into lipid nanoparticles (LNPs) specially formulated to deliver RNAi into primary mouse neurons using the Neuro9 siRNA Spark Kit (Precision Nanosystems). Cells were transfected with 0.3 μg/ml of siRNA/GAPmer supplemented with 1μg/ml ApoE4 on DIV 7. A fluorescent control siRNA was used to confirm a transfection efficiency of more than 80%. Knockdown efficiency was initially validated by qPCR after 48 hours but sufficient decrease in Mettl3 protein and m^6^A levels were reached with the use of GAPmers for 6 days after transfection. Before fixation or RNA or protein extraction cells were washed with sterile DPBS to remove medium.

### m^6^A quantification

m^6^A concentration was determined using a m^6^A Methylation Assay Kit Fluorometric (Abcam). The starting material was 200 ng of rRNA-depleted RNA and the manufacturer’s protocol was followed. All reactions were carried out in duplicate and a standard curve of m^6^A /A was included to have quantitative results. Reactions were read in a FLUOstar Omega Multiplate reader (BMG) in fluorescence mode.

### Western blot/Immunofluorescence

Antibodies used for Western blot, immunofluorescence and other applications, as well as the dilutions used are described in Supplementary Table 2.

### Puro-PLA

Puromycin-proximity ligation assay (Puro-PLA) was performed as previously described with minor alterations ^50^. DIV 13 mouse primary hippocampal neurons were pretreated with 100 ug/ml cycloheximide for 30 minutes to arrest translational elongation. Cells were then treated with 3 μM puromycin for 10 minutes to label nascent polypeptide chains. This treatment time was chosen to balance labeling intensity with the propensity of labeled peptides to diffuse away from their synthesis sites ^51,90^. Puromycin incorporation and cycloheximide pretreatment were validated by Western blot. The PLA was performed using the Duolink Proximity Ligation Assay Kit Red (Merck) according to the manufacturer’s instructions. Primary antibodies against puromycin and the protein of interest (targeting the N-terminal region) raised in different species were used for the PLA assay and incubated ON at 4°C. Counterstain antibodies (Map2 and SYP) were incubated ON at 4°C along with the PLA primary antibodies, an additional incubation step was added after finishing the Puro-PLA protocol to add the secondary antibodies for the counterstains for 2 hours at RT before mounting.

For each condition and replicate, 7-13 neurons were captured by confocal imaging and analyzed using Cell Profiler to automate the analysis and remove biases. Background level PLA signal was adjusted to samples without puromycin treatment.

### Polysome sequencing

Polysomes were prepared from the DG of five young and five old animals as described ^91^.

### Synaptosome isolation for sequencing

Synaptosomes were isolated from the hippocampi of 3-month and 16 months old as recently described ^38^.

### H3K36me3 ChIP

Cell type specific chromatin isolation and ChIP sequencing was performed as previously described ^92 93^. 3-4 CA1 were pooled for each replicate and nuclei were FACS sorted by NeuN expression. 300ng of chromatin and 1μg of H3K36me3 antibody (Abcam, ab9050) were used for each ChIP.

## Data availability

GEO database: GSE198526. For review, the token can be obtained via the editor.

## Acknowledgments

This work was supported by the following third-party funds to AF: The DFG priority program 1738, SFB1286, EPIFUS project. MEB and MTB were supported by the DFG priority programme 1784. AF and MTB were supported by Germany’s Excellence Strategy - EXC 2067/1 390729940.

## Notes

### Competing Interest Statement

The authors have declared no competing interest.

