## Supplementary figures for "Conserved reduction of m^6^A marks during aging and neurodegeneration is linked to altered translation of synaptic transcripts"

**Supplemental Figure 1**

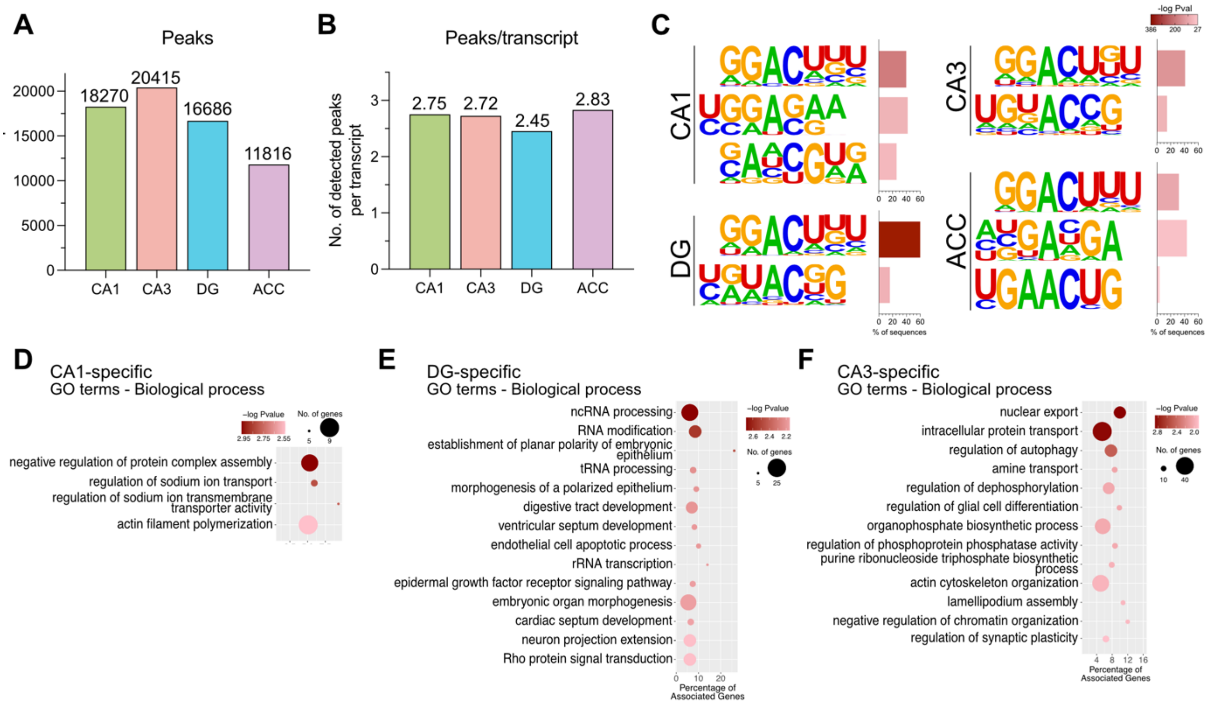


**Figure S1. m$^6$A levels in the brain of 3 months old mice A.** Detected m$^6$A peaks in hippocampal subregions and the ACC in the adult mouse brain. **B.** Average number of m$^6$A peaks detected per methylated transcript. Exact values are displayed above the corresponding bar. **C.** Enriched motifs corresponding to the DRACH consensus sequence among the top overrepresented motifs in m$^6$A peaks. **D-F.** Enriched GO terms Biological processes for CA1- (D), DG- (E) and CA3-specific (F) methylated transcripts.

**Supplemental Figure 2**

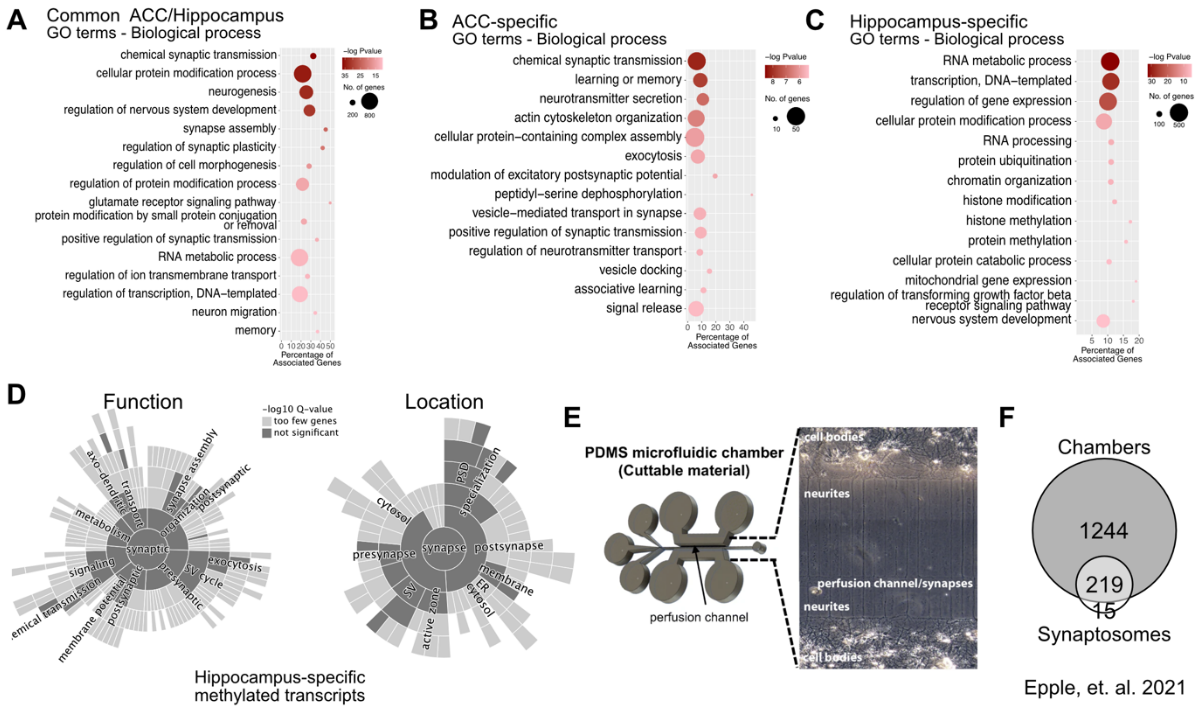


**Figure S2. Characteristics of the m$^6$A transcripts detected in the brain of 3 months old mice. A-C.** Enriched GO terms biological process for methylated transcripts common between the hippocampus and the ACC (A), specific for the ACC (B) and specific for the hippocampus (C). **D.** Synaptic GO showing the lack of significant enrichment for synaptically annotated Function or Location GO terms in hippocampus-specific methylated transcripts**. E.** Diagram showing the microfluidic chambers used by Epple, et. al (2021) to isolate synaptically located RNAs. **F.** Summary of the synaptic transcriptome as described by {Epple, 2021}.

**Supplemental Figure 3**

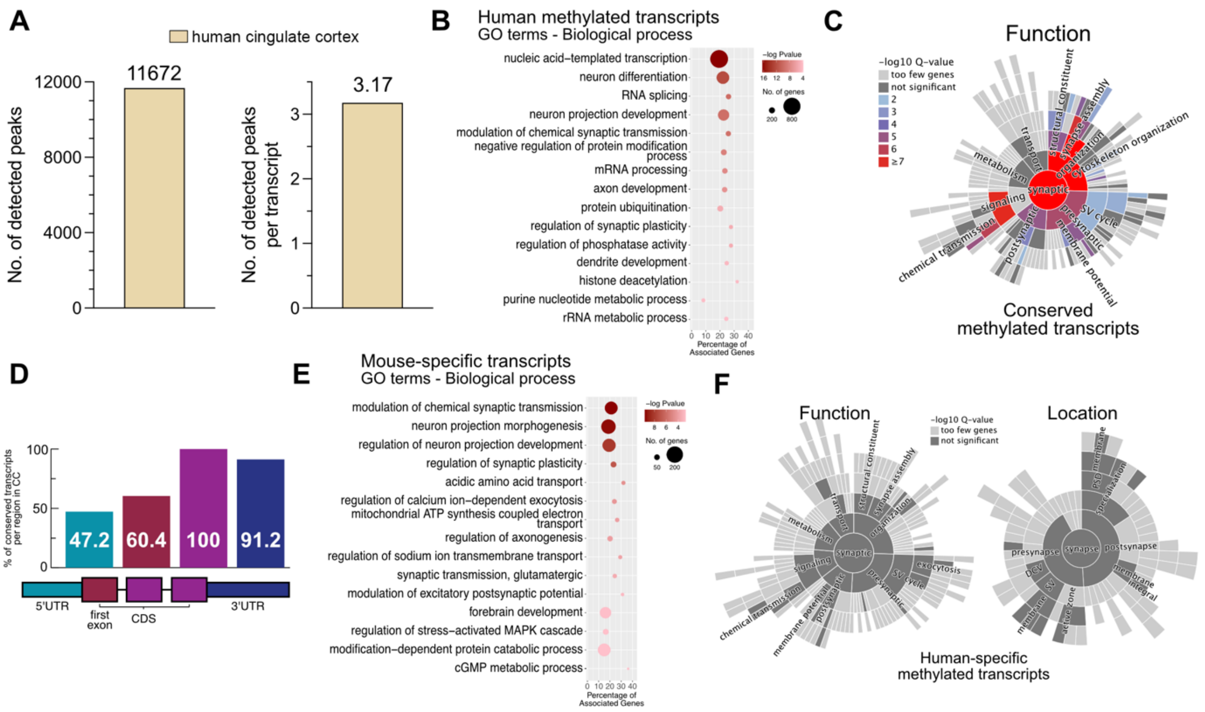


**Figure S3. Characteristics of the m$^6$A transcripts detected in the human cingulate cortex. A**. Number of detected m$^6$A peaks (left panel) and average number of peaks per methylated transcript (right panel) in the human CC. Exact number are displayed above the corresponding bar. **B.** Enriched GO terms Biological process for all methylated transcripts in the human CC. **C.** Synaptic function annotated GO terms enriched in methylated transcripts conserved in mouse/human. **D.** Regional conservation of methylation sites in mouse/human. Bars show the percentage of peaks annotated for a given region in the human CC that have a corresponding peak annotated to the same region in the mouse homolog. Only broad regions were considered: 5'UTR, first exon, CDS and 3'UTR. **E.** Enriched GO terms Biological process for mouse specific methylated transcripts from the ACC. **F**. Synaptic GO terms for function and localization showing the lack of significant enrichment in these categories of human-specific methylated transcripts.

**Supplemental Figure 4**

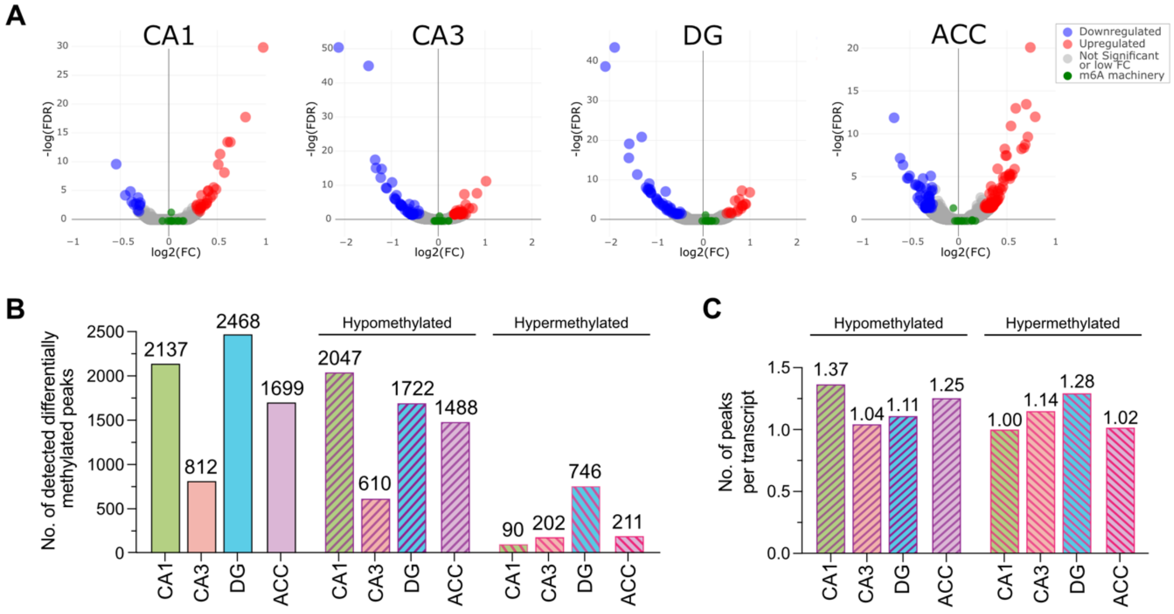


**Figure S4. Changes in gene-expression and m$^6$A peaks in the aging mouse brain. A.** Volcano plots displaying the changes in gene expression across brain subregions comparing 3 vs 16 months old mice. Cutoffs for significance are FC > 1.2 and FDR ≤ 0.05. Highlighted in green are the known m$^6$A writers, readers and erasers, showing that no m$^6$A associated protein is differentially expressed in the aged brain. **B.** Bar graphs showing the total amount differentially methylated peaks (FC > 1.2, FDR ≤ 0.05) in the brain subregions and how many of them are hypo- and hypermethylated. **C.** Bar graphs showing the number of m$^6$A peaks per differentially methylated transcript. Numbers above the bars display the exact number of peaks.

**Supplemental Figure 5**

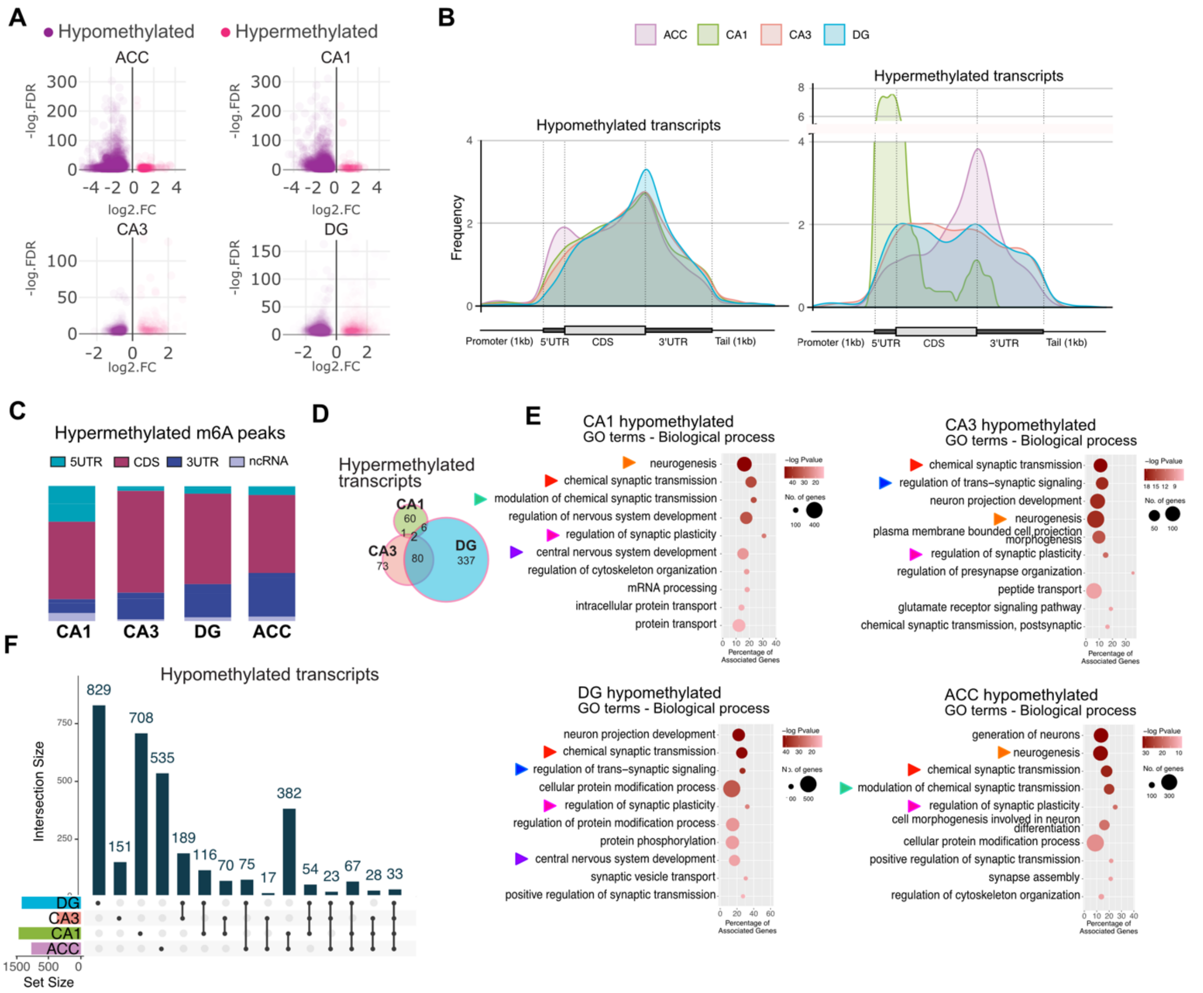


**Figure S5. m$^6$A levels in the aging mouse brain. A**. Volcano plots showing the magnitude of changes in m$^6$A for every differentially methylated peak in brain subregions comparing 3 vs 18 months old mice. **B.** Guitar plots showing the frequency of hypomethylated (left panel) and hypermethylated (right panel) peaks along mRNA features in all investigated brain regions. **C.** Distribution of hypermethylated peaks along RNAs in all brain subregions. **D.** Overlap of hypermethylated transcripts across hippocampal subregions. **E.** GO terms biological process for transcripts hypomethylated specifically within the indicated brain region. **F.** Intersect graph showing the overlap between hypomethylated transcripts across all brain subregions, as well as the region-specific hypomethylated transcripts. Dots and lines denote the displayed comparison and bars show the number of transcripts contained in the overlap. C. ACC - anterior cingulate cortex, DG - dentate gyrus, 5UTR - 5' untranslated region, 3UTR - 3' untranslated region, CDS - coding sequence, ncRNA - non-coding RNA.

**Supplemental Figure 6**

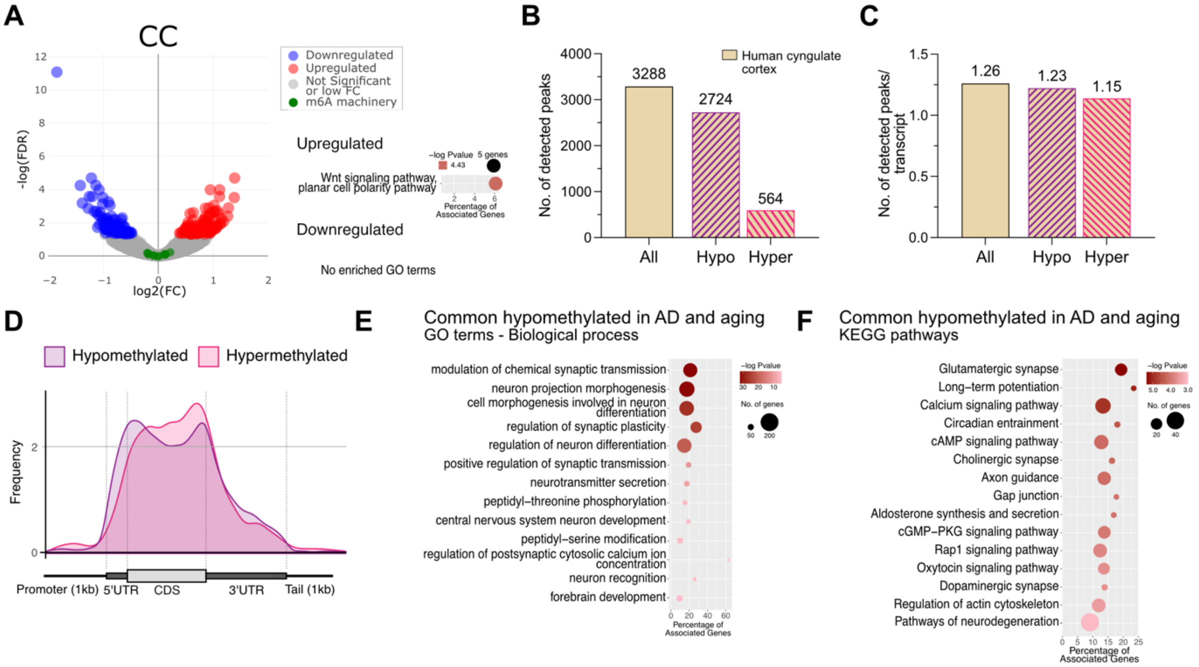


**Figure S6. Altered m$^6$A levels in in the cortex of human AD patients. A.** volcano plot displaying the changes in gene expression in the human CC in AD (FC > 1.2, FDR ≤ 0.05). Highlighted in green are the known m$^6$A writers, readers and erasers, showing their unaltered expression. Right panel shows significantly enriched GO term Biological process for upregulated genes. Downregulated genes resulted in no enriched GO terms. **B.** Total number of differentially methylated m$^6$A peaks in AD compared to control samples, as well as hypo- and hypermethylated peaks. **C.** Average detected differentially methylated m$^6$A peaks per differentially methylated transcript in AD. **D.** Guitar plot showing the distribution frequency of hypo- and hypermethylated peaks along mRNA features. **E, F.** Enriched GO terms Biological process (E) and enriched KEGG pathways (F) for commonly hypomethylated transcripts from all peaks in AD and the aged ACC in mouse (converted to their corresponding human homolog). CC - cingulate cortex, 5'UTR - 5' untranslated region, 3'UTR - 3' untranslated region, CDS - coding sequence, ncRNA - non-coding RNA

**Supplemental Figure 7**

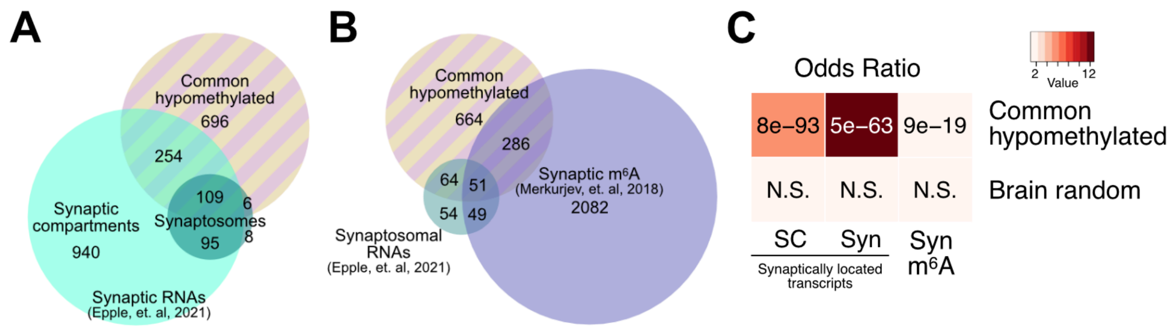


**Figure S7. Transcripts hypomethylated in the aging mouse and human Ad brain are enriched at synapses. A.** Venn diagram showing the overlap between commonly hypomethylated transcripts shown in main Fig 4E and synaptically located RNAs detected in hippocampal synaptosomes or synaptodendritic compartments of neuorns cultured in microfluidic chambers as obtained from Epple, et. al. 2021. **B.** Overlap between commonly hypomethylated transcripts, synaptosomal RNAs from Epple, et. al, 2021 as well as synaptic (syn) m$^6$A transcripts from Merkurjev, et. al, 2018. **C.** Enrichment (odds ratio) and significance of the overlaps displayed in A and B. Color represents odds ratio of the comparison, p value is visible in the corresponding square. “Brain random” corresponds to a list of 2000 randomly selected brain-expressed transcripts, included as control. SC – synaptic compartments, Syn – synaptosomes, Syn m6A – synaptic m6A mRNAs, N.S. – not significant.

**Supplemental Figure 8**

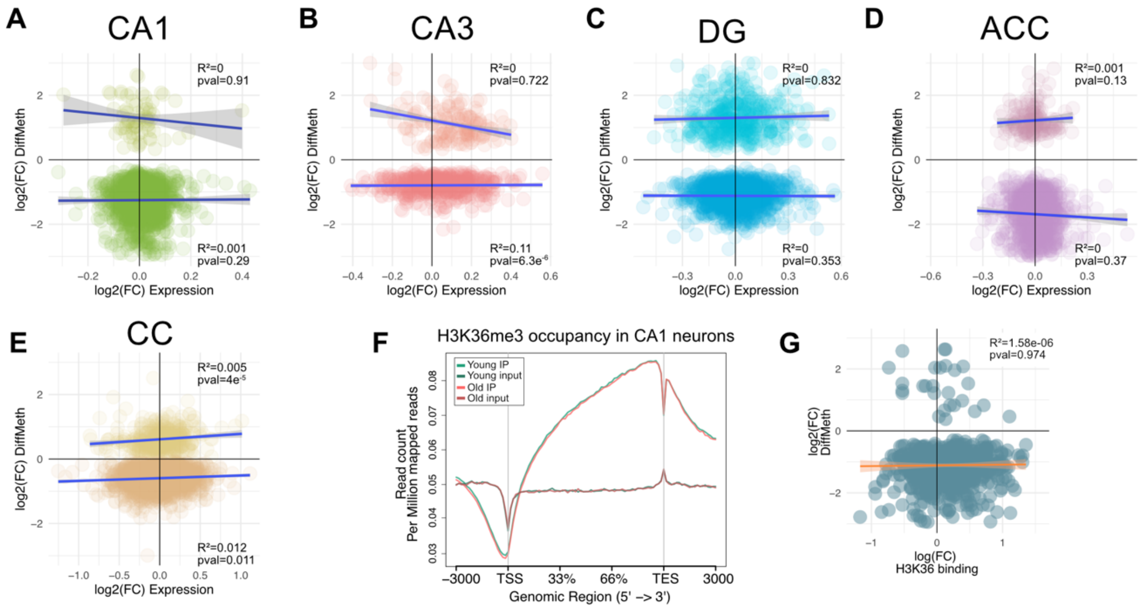


**Figure S8. Changes in N$^6$A RNA-methylation are not linked to altered mRNA levels or H3K36me3. A-E.** Scatter plots showing the correlation between observed changes in m$^6$A RNA-methylation and changes in transcript levels in the CA1 (A), CA3 (B), DG (C), and ACC (D) regions of 3 vs 16 months old mice as well as in the human CC (E) of control vs AD patients. **F.** Occupancy of H3K36me3 in FACS sorted CA1 neuronal nuclei determined by ChIP. Curves show normalized reads mapped to the shown genomic regions in immunoprecipitated (IP) and input samples from young and old mice. **G.** Scatter plot displaying changes in H3K36me3 occupancy and m$^6$A changes in all differentially methylated transcripts in the aged CA1. In all scatter plots, line represents the best model fit for the data points, CI is also displayed. Reads in CPM. TSS = transcription start site, TES = transcription end site.

**Supplemental Figure 9**

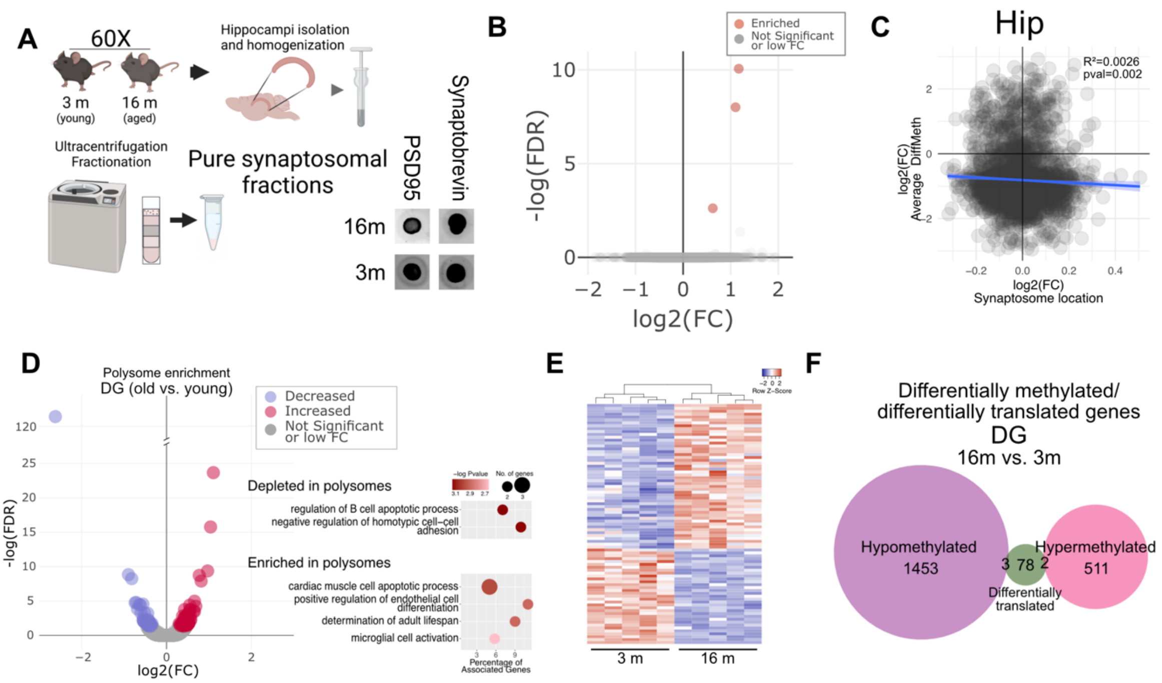


**Figure S9. Synaptosomal and polysomal RNA in the young vs aged brain. A** Diagram showing synaptosomal fraction purification from the hippocampus of young (3 month) and old mice (16 month) as described previously {Epple, 2021}. Synaptosome purity was confirmed by dot blot using the synaptic markers PSD95 and synaptobrevin. **B.** Volcano plot showing differential expressed synaptosomal RNAs in the old vs. young hippocampus. Only three genes were significantly different. **C.** Scatter plot showing that there is no obvious correlation between the expression level of synaptosomal transcripts and their methylation level **D.** Volcano plot displaying the results of a differential expression analysis performed on polysomes isolated from the hippocampal DG region of young (3 month) and old mice (16 month). GO terms Biological process overrepresented in mRNAs enriched and depleted in polysomal fractions are shown (right panel). **E.** Heatmap displaying the differentially translated mRNAs across replicate polysome samples. **F.** Venn diagram of differentially translated and differentially methylated mRNAs in the aged DG showing little overlap between these populations.

**Supplemental Figure 10**

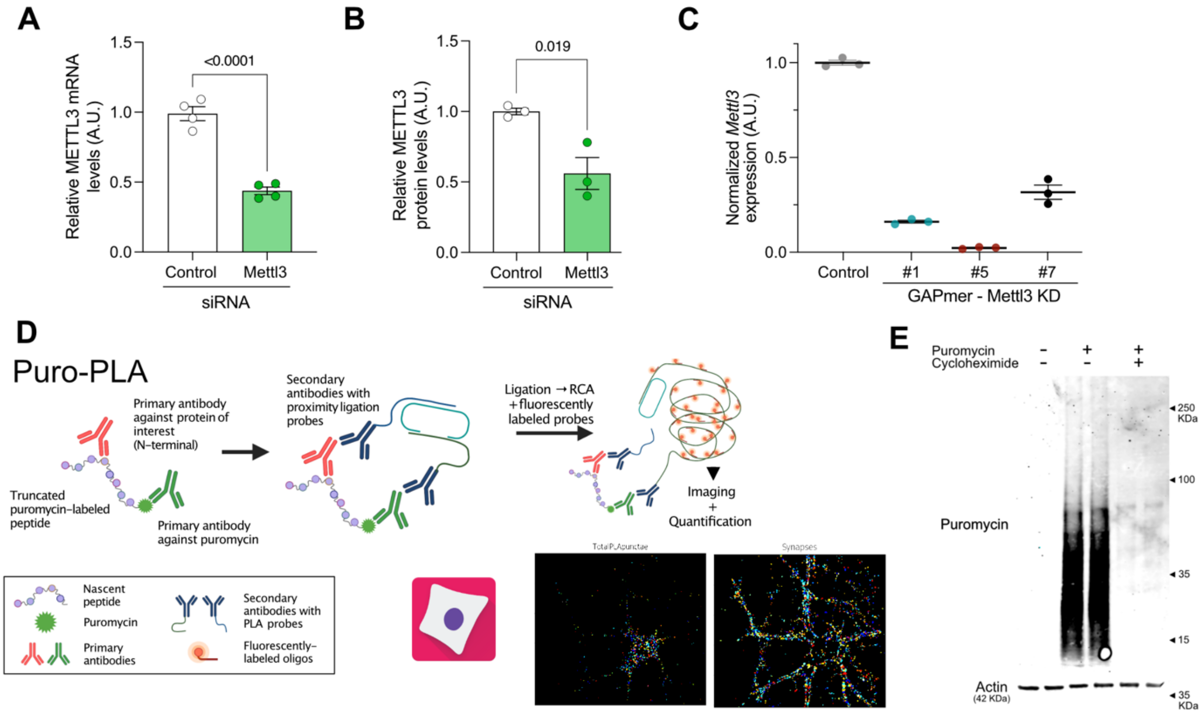


**Figure S10. Knock down of *Mettl3* via GAPmers to study local protein synthesis A.** Bar graph showing that *Mettl3* knock-down via siRNAs partially decreases *Mettl3* mRNA at and protein levels **(B)** levels in hippocampal neurons after 48 hours of incubation. **C.** Validation of LNA GAPmer-dependent knock-down of *Mettl3* in primary hippocampal neurons. Shown are the mRNA levels of Mettl3 after 48 hours treatment with control or Mettl3-targeting GAPmers (1,5 and 7). GAPmer #5 was selected for further experiments. Graphs in A-C display the mean +/- SEM of each condition. Each data point represents one independent replicate, statistical significance was determined by Student's t test. **D.** Scheme of the Puro-PLA process. Shown are representative images from the automated Puro-PLA and synapse detection pipeline using Cell Profiler. **E.** Validation of puromycin incorporation into nascent protein chains. Western blot using a puromycin antibody showing the labeling of proteins and the function of the cycloheximide pretreatment to cause translational arrest.

**Supplemental Figure 11**

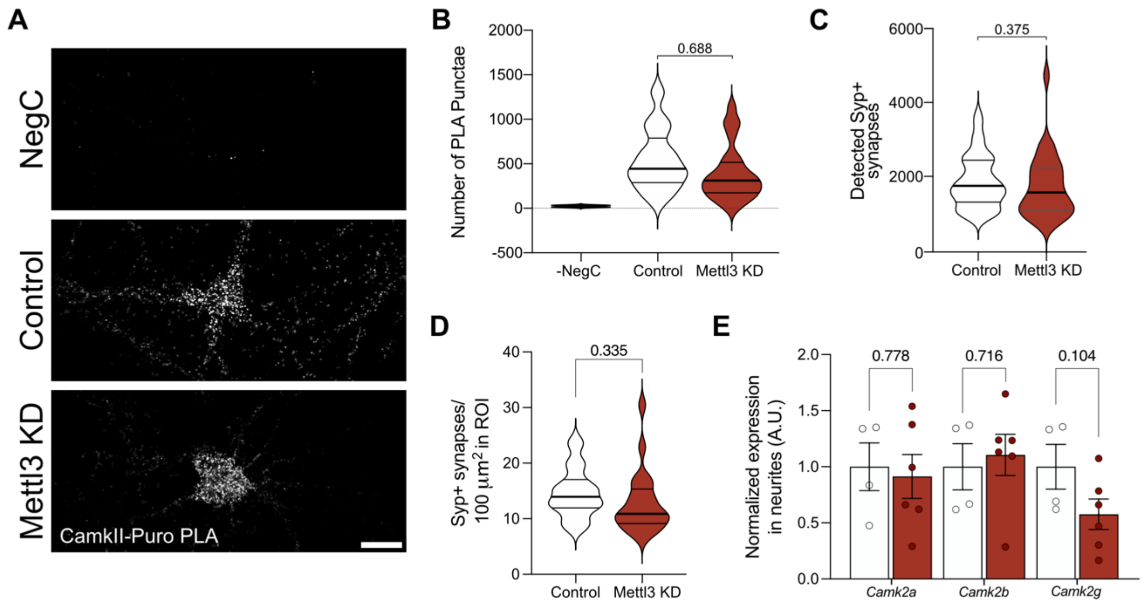


**Figure S11. Mettl2 knock-down does not affect total amount of CAMKII production, synapse number or synaptic elvels of CamKII transcripts. A.** Representative confocal images from untreated (without puromycin treatment, Puromycin-), control and Mettl3 GAPmer-treated cells showing CAMKII-Puro PLA punctae. Scale bar: 10µm. **B.** Violin plots showing Total number of detected CAMKII-Puro PLA punctae in Puromycin-, control and *Mettl3* KD neurons. **C, D.** Total number (C) and area normalized number (D) of synaptophysin+ (SYP+) synapses in Control and Mettl3 KD neurons. Graphs in B, C and D show the mean of 3 independent experiments, for each experiment 7-13 neurons were imaged and analyzed, individual data points were used to generate the violin plot. Quartiles are marked by gray lines. Statistical significance was determined by Student's t test on the mean values of each independent replicate. **E.** mRNA levels, determined by qPCR, of different CAMKII isoforms in the synaptic compartments of microfluidic chambers containing neurons treated with Control or Mettl3 GAPmers. Graphs display the mean +/- SEM of each condition. Each data point represents one independent replicate, statistical significance was determined by Student's t test
